# A Comparative Chemical-Genetic Screen Reveals Divergent Roles for Host ALK2 Signaling Across Intracellular Bacterial Niches

**DOI:** 10.64898/2026.07.30.741746

**Authors:** Sahoo Parnika, Malavika K A, Mullai Valli Ramamoorthy, Usha Krishna, Aishani Saha, Megha Manoj Menon, Ling Wei, Sandhya Ganesan, Kamalakannan Vijayan

## Abstract

Diverse vacuolar architectures present unique signaling challenges for intracellular bacteria, yet whether distinct niches share host requirements remains unclear. Because host kinases coordinate these complex cellular responses, we performed a comparative chemical-genetic screen of *Coxiella burnetii*, *Chlamydia trachomatis*, and *Salmonella enterica*. The screen revealed that *C. burnetii* is uniquely dependent on host kinase pathways. Deconvolution of these inhibitor profiles identified the bone morphogenetic protein (BMP) type I receptor, ALK2, as a critical regulatory checkpoint. While *C. burnetii* activates ALK2-mediated phospho-SMAD1/5/9 signaling to support replication, *C. trachomatis* actively suppresses this cascade. Mechanistically, ALK2 functions upstream of TFEB nuclear translocation and LAMP1 recruitment, driving the lysosomal biogenesis required to sustain the phagolysosome-like niche of *C. burnetii* while disrupting the non-lysosomal inclusion of *C. trachomatis*. Consequently, targeting ALK2 switches replication outcomes, demonstrating how distinct niches implement opposing strategies to exploit a single host signaling node.

## Introduction

Obligate intracellular bacterial pathogens must engineer a specialized vacuole that shields them from host cytosolic immune sensors. However, such compartmentalization inadvertently creates a nutrient barrier, forcing the pathogens to remodel the vacuolar membrane to import essential host nutrients ^1–4^. To overcome this protection-versus-starvation paradox, these pathogens actively remodel host signaling pathways to acquire nutrients and maintain their intracellular niche. For example, *Coxiella burnetii* enters through the endocytic pathway to establish a highly acidic *Coxiella*-containing vacuole (CCV) that relies on continuous organelle fusion, particularly with lysosomes ^5,6^. In contrast, *Chlamydia trachomatis* generates a non-fusogenic, neutral-pH inclusion that segregates from the endolysosomal network and intercepts exocytic nutrient trafficking ^7,8^, while *Salmonella enterica* maintains a dynamic *Salmonella*-containing vacuole (SCV) characterized by extensive membrane tubulation events that facilitate interactions with host trafficking pathways^9^. Whether these diverse intracellular pathogens rely on distinct host signaling programs or converge on a common infection-permissive cellular state remains poorly understood.

Host kinase signaling networks represent central regulatory hubs that coordinate virtually all aspects of cellular physiology. As such, they constitute vulnerable control points that intracellular pathogens can precisely access and reprogram to balance vacuolar integrity with nutrient acquisition. *C. burnetii* promotes survival within its metabolically demanding vacuole by activating host pro-survival signaling pathways including AKT and ERK1/2 ^10,11^, while *C. trachomatis* recruits host lipid, signaling, and trafficking kinases to coordinate inclusion expansion and intracellular development ^12,13^.

Early chemical-genetic studies attempted to map these dependencies by treating host phosphorylation cascades as isolated, single-target pathways. For example, screening a targeted panel of kinase inhibitors during *C. burnetii* infection identified host requirements for protein kinase C (PKC), p38 MAPK, and cAMP-dependent protein kinase (PKA) signaling during vacuolar maturation ^14^, while similar studies in *C. trachomatis* identified host signaling pathways associated with inclusion development ^15^. Although these foundational studies established that host signaling networks contribute to vacuole maintenance, their mechanistic interpretation was limited by reliance on nominal drug-target annotations. By assuming that kinase inhibitors interact exclusively with their intended targets, these approaches overlooked widespread kinome-wide cross-reactivity and off-target activities, leaving the true causal signaling networks unresolved. Uncovering which host kinase nodes represent universally conserved master regulators versus niche-specific drivers requires bypassing nominal annotations, a challenge we address here using integrated phenotypic-polypharmacological deconvolution.

Rather than treating multi-target cross-reactivity as a confounding experimental limitation, contemporary chemical biology frameworks leverage polypharmacology as an information-rich signal for target deconvolution ^16^. This computational strategy, originally developed by Gujral and colleagues to mathematically resolve phenotypic target identities from multi-kinase activity profiles ^17,18^, was subsequently deployed to identify host regulators of *Plasmodium* liver-stage infection ^19^. More recently, machine learning-assisted regularization approaches have been used to dissect complex physiological networks such as endothelial barrier regulation ^20^ and to define host kinase architectures governing intracellular protozoan infection ^21^. By regressing phenotypic screening data against experimentally determined inhibitor-kinase activity matrices using penalized regression models, these approaches transform overlapping inhibitor selectivity profiles into a quantitative map of causal signaling dependencies. Applying this systems-level framework to intracellular bacterial pathogens provides an opportunity to identify both conserved and pathogen-specific host signaling architectures that cannot be resolved through conventional inhibitor screening approaches.

To systematically define these pathways, we performed a comparative chemical-genetic deconvolution analysis across *C. burnetii, C. trachomatis*, and *S. enterica*. By integrating multi-pathogen phenotypic screening data with machine learning-based regression of inhibitor-kinase activity profiles, we mapped both conserved and pathogen-specific host kinase dependencies. This approach identified the bone morphogenetic protein (BMP) type I receptor ALK2 as a candidate host signaling node shared across these distinct intracellular niches. We subsequently evaluated the role of ALK2 signaling and downstream SMAD activation using complementary genetic and pharmacological approaches and further leveraged the predictive framework to perform virtual screening for host-directed inhibitors. Together, these findings establish a systems-level framework for resolving host kinase vulnerabilities and reveal a previously unrecognized role for ALK2 signaling during intracellular bacterial infection.

## Results

### A Multi-Pathogen Chemical Screen Uncovers Divergent Host Kinase Vulnerabilities Across Vacuolar Niches

To systematically map host signaling pathways exploited by vacuolar bacterial pathogens, we performed a comparative chemical-genetic screen using a library of 38 small-molecule kinase inhibitors. Three distinct intracellular niches, the acidic *Coxiella*-containing vacuole (CCV), the neutral *Chlamydia* inclusion, and the dynamic *Salmonella*-containing vacuole (SCV), were subjected to an identical chemical library in human epithelial (HeLa) cells (**Figure 1A and B**). Infection frequency (% infection) and intracellular replication (% development) were quantified in parallel, while host cell viability (**Figure S1A**) was monitored to exclude cytotoxicity-driven artefacts.

**Figure 1.**
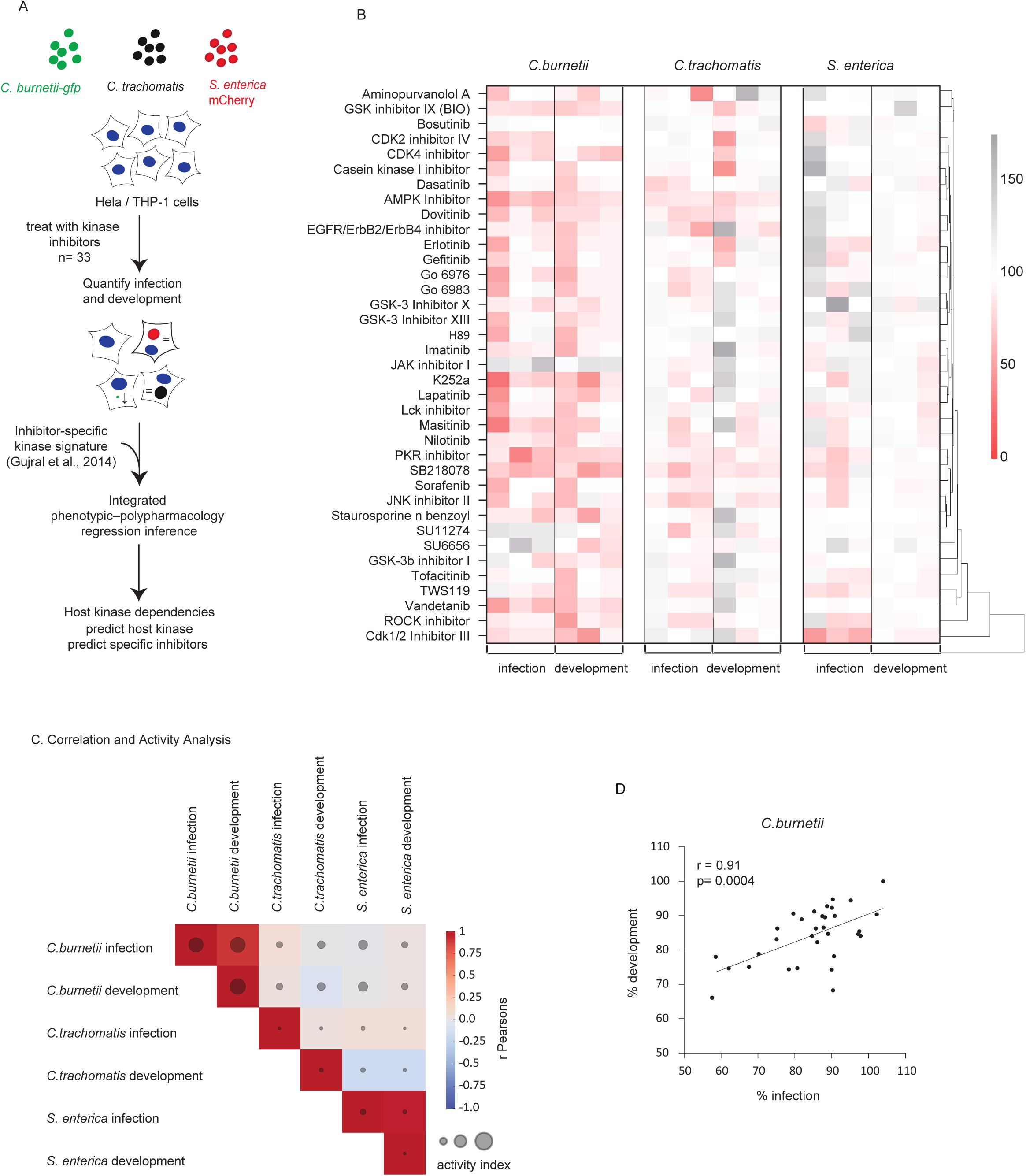
Comparative chemical-genetic screen reveals divergent host kinase dependencies across intracellular vacuolar niches. (**A**) Schematic of the screening workflow of cells infected with *Coxiella burnetii* GFP, *Chlamydia trachomatis*, or *Salmonella enterica-* mCherry, followed by treatment with a panel of kinase inhibitors and quantification of infection frequency and intracellular development; inhibitor-specific kinase activity profiles were then used for regression-based target deconvolution. (**B**) Heatmap showing the normalized effects of kinase inhibitors on the infection frequency and development across the three infection systems. (**C**) Pearson correlation and activity index analysis across all screening outputs. (**D**) Strong positive correlation between *C. burnetii* infection and development phenotypes (r = 0.91, p = 0.0004).

The screen revealed highly divergent, pathogen-specific dependencies across the tested chemical space (**Figure 1C**). *C. burnetii* displayed a profound and widespread vulnerability to kinome perturbation, with multiple inhibitor chemotypes markedly reducing both infection and intracellular development. To confirm that these phenotype reductions reflected host-directed mechanisms rather than off-target antibacterial toxicity, we evaluated the inhibitor panel against *C. burnetii* in host-free axenic growth media. None of the compounds impaired bacterial growth at the screening concentrations (**Figure S1B**), establishing that the observed anti-infective effects were strictly host-mediated. In contrast, *C. trachomatis* exhibited a more restricted susceptibility profile, while *S. enterica* showed minimal phenotypic sensitivity to the inhibitor library. Together, these results reveal a clear hierarchy in host kinase dependence, with *C. burnetii* displaying the greatest reliance on host phosphorylation networks among the pathogens examined.

To determine whether early host cell colonization and downstream vacuolar development require shared or distinct signaling pathways, we performed Pearson correlation and activity index analyses across all screening outputs (**Figure 1C**). While inter-pathogen phenotypic correlations were generally weak, strong intra-pathogen relationships were observed. This coupling was most pronounced for *C. burnetii*, where infection and development exhibited a strong positive correlation (r = 0.91, p = 0.0004). These findings suggest that host kinase perturbations that disrupt early *C. burnetii* niche establishment similarly influence downstream replication, providing a robust dataset for computational target deconvolution.

### Secondary Screening in Macrophages and Polypharmacological Deconvolution Uncover Cell-Type-Specific and Shared Host Targets

To evaluate whether the identified host kinase dependencies are cell-line dependent and to validate these vulnerabilities in a more physiologically relevant niche, we performed a secondary chemical screen on infected THP-1 human macrophage-like cells (**Figure 2A**). This library-wide secondary evaluation demonstrated a broad spectrum of intracellular burdens, revealing that specific small-molecule chemotypes, such as Cdk1/2 Inhibitor III, K252a, Dasatinib, and an AMPK inhibitor, profoundly reduced the total *C. burnetii* burden within macrophages. Intersecting this macrophage screening data with the primary invasion and development datasets from HeLa cells, we mapped a robust core signature of consensus hits that restrict *C. burnetii* pathogenesis across multiple diverse cellular environments. To bridge the gap between small-molecule sensitivity and discrete molecular targets, we leveraged published quantitative biochemical selectivity matrices to evaluate the target profiles of our active compounds. Recreating these kinase activity profiles as a residual activity matrix highlighted a select group of candidate kinases that are significantly and potently inhibited by these active chemotypes (**Figure 2B**).

**Figure 2.**
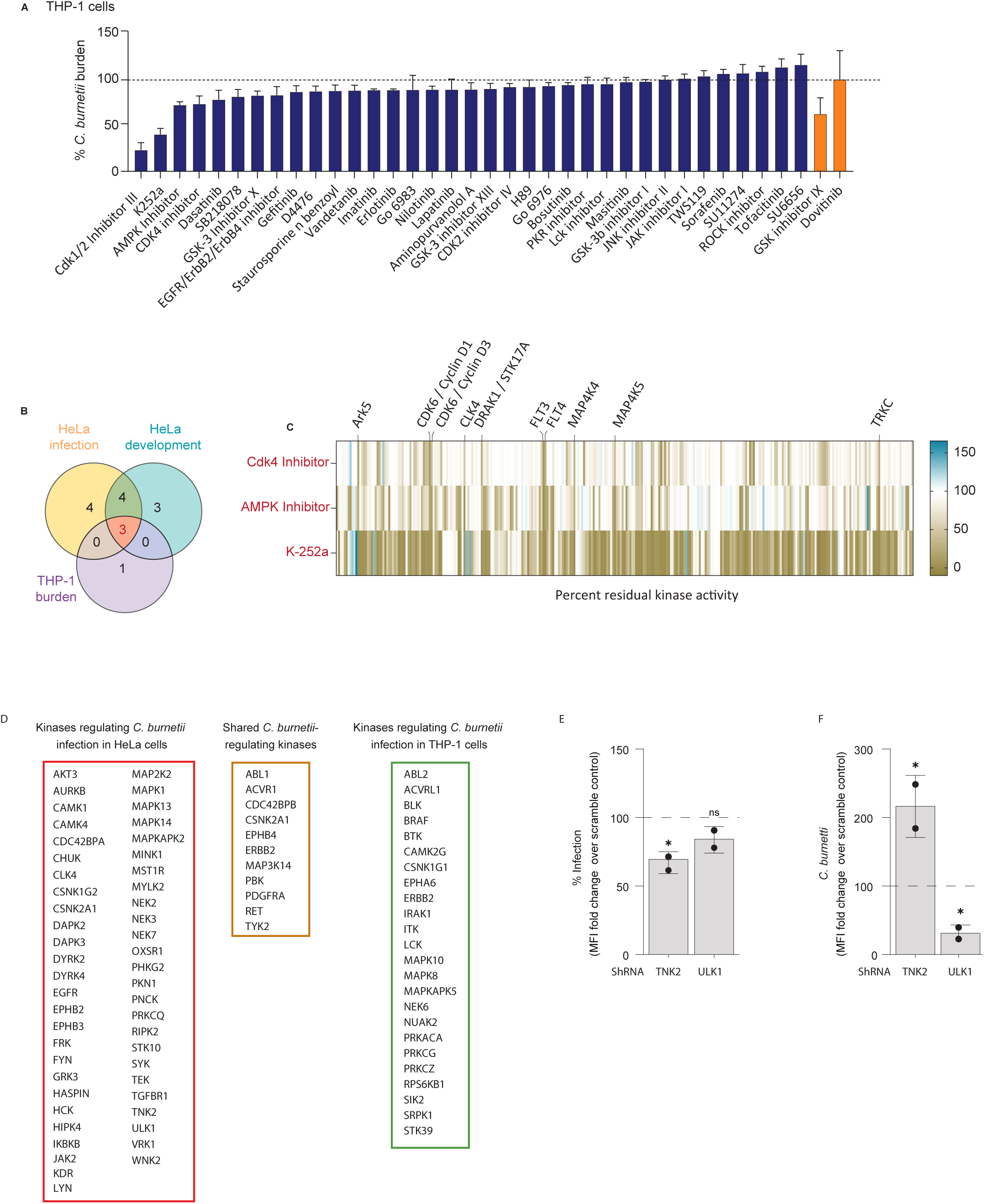
Polypharmacology-based deconvolution identifies shared and cell-type-specific host kinase dependencies regulating *C. burnetii* infection. (**A**) Secondary kinase inhibitor screen in THP-1 macrophages infected with *C. burnetii* identifying compounds that alter intracellular bacterial burden. The dashed line indicates the DMSO control. (**B**) Overlap analysis of kinase inhibitor hits identified from HeLa infection, HeLa development, and THP-1 burden datasets. (**C**) Residual kinase activity profiles of representative active compounds (CDK4 inhibitor, AMPK inhibitor, and K252a) used for target deconvolution. (**D**) Elastic net regression identifies host kinases associated with *C. burnetii* infection in HeLa cells, THP-1 macrophages, and a shared set of conserved regulators across both cellular contexts. (**E, F**) Functional validation of predicted host regulators by shRNA-mediated knockdown of TNK2 and ULK1, followed by quantification of infection burden (**E**) and *Coxiella*-containing vacuole (CCV) MFI (mean fluorescence intensity) (**F**). Data are presented as fold change relative to scramble shRNA controls. *P < 0.05; ns, not significant.

To translate these overlapping polypharmacological fingerprints into a predicted map of molecular targets, we applied an elastic net regularized regression model to deconvolute the primary and secondary kinome data ^22^. This systems-level mathematical model successfully partitioned the host kinome into distinct cell-type-specific vulnerabilities and universally conserved nodes (**Figure 2C**). Specifically, the model isolated a cluster of host kinases (**Figure 2D**) uniquely required within the HeLa epithelial niche (including AKT3, AURKB, MAP2K2, and MAPK1) and an independent set of kinases restricted to the THP-1 macrophage-like environment (such as ABL2, ALK2, MAPK13, CDC42BPB, MAPK14, CSNK2A1, EGFR, and ERBB2). Crucially, the deconvolution resolved a core set of shared regulatory nodes, most notably the non-receptor tyrosine kinase ABL1 and the bone morphogenetic protein (BMP) type I receptor, ALK2, that are fundamentally required to support infection across both host cell architectures (**Figure 2D**).

To experimentally validate these computational predictions *in vitro*, we performed targeted lentiviral shRNA-mediated knockdown of specific prioritized nodes (**Figure S2**), monitoring both colonization rates and physical vacuolar morphology. In precise agreement with the model’s predictive framework, genetic silencing of the predicted promoter ULK1 severely compromised intracellular pathogenesis, significantly reducing overall infection rates and arresting *Coxiella*-containing vacuole (CCV) expansion relative to scramble controls (**Figure 2E, 2F**). Conversely, depletion of the predicted restrictor TNK2 induced a robust hyper-infection phenotype, driving a dramatic increase in both the percentage of infected host cells (∼150%) and final CCV geometric dimensions (∼200-300%) (**Figure 2E, 2F**). Standard annotation-based screening analyses would have merely highlighted the intended primary targets of these compounds, failing to resolve these non-canonical drivers. Collectively, these findings demonstrate that bypassing nominal drug annotations through regularized deconvolution accurately uncovers both niche-specific and universally conserved host signaling nodes regulating vacuolar maintenance.

### Global Kinome Mapping and Pathway Enrichment Analysis Define the Regulatory Host Landscape

To evaluate the host kinase networks driving or restricting *C. burnetii* pathogenesis, we first segregated our regularized elastic net deconvolution hits based on their cellular niche and phenotypic directionality (**Figures 3A and 3B**). Next, we globally visualized these targets by mapping them across the human kinome phylogenetic tree (**Figure 3C**). This systems-level topological analysis predicted a robust network of unique kinases that support infection and that actively suppress it.

**Figure 3.**
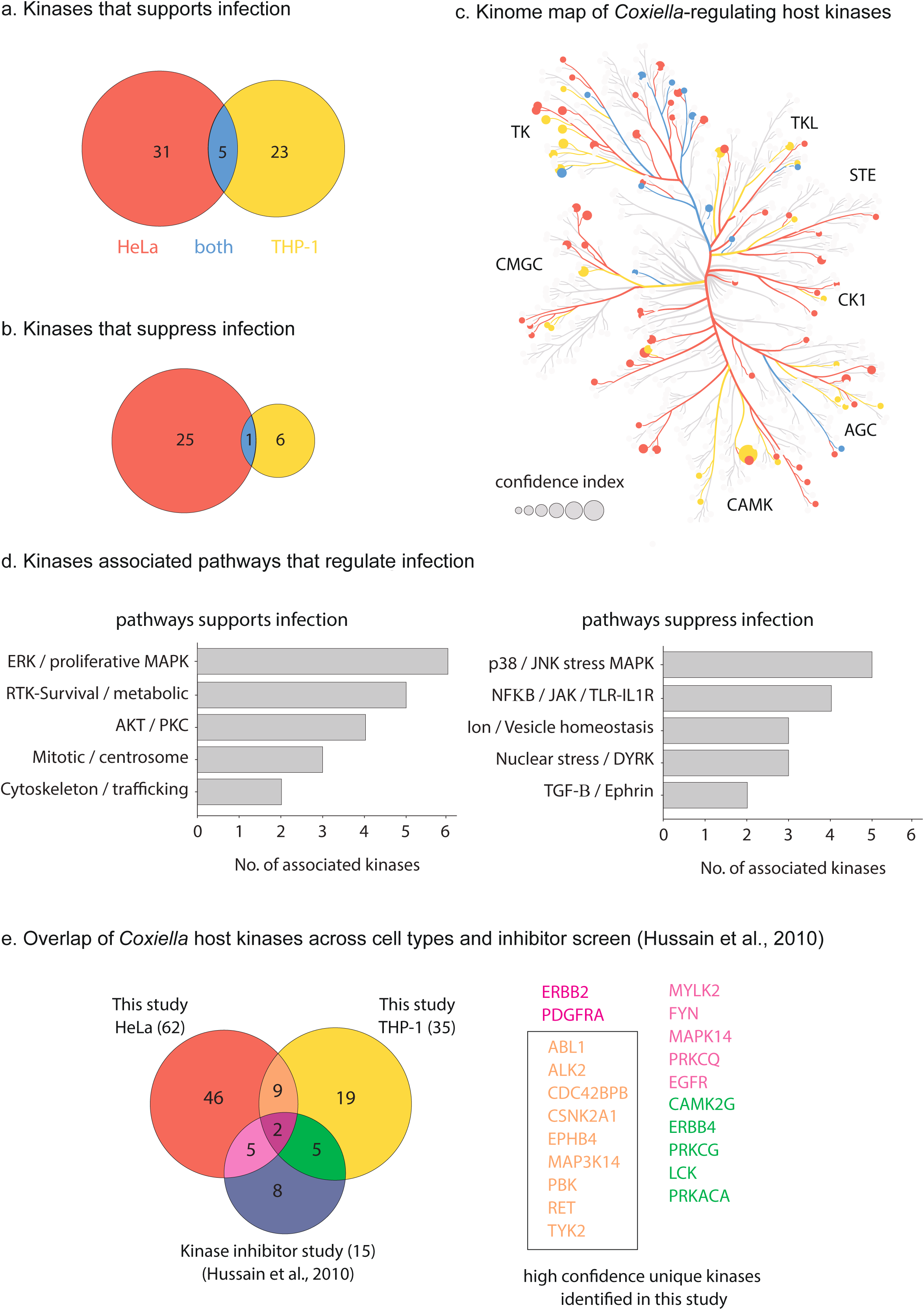
Global kinome mapping reveals host signaling pathways associated with *C. burnetii* infection. (**A**) Overlap of host kinases predicted to support *C. burnetii* infection in HeLa epithelial cells and THP-1 macrophages. (**B**) Overlap of host kinases predicted to suppress *C. burnetii* infection across both cellular contexts. (**C**) Distribution of *C. burnetii*-associated host kinases across the human kinome phylogenetic tree. Node size indicates confidence index, and colors denote kinase family classification. (**D**) Functional classification of infection-associated and development-associated kinases, highlighting enrichment of receptor signaling, MAPK-driven growth, metabolic adaptation, stress adaptation, cytoskeletal remodeling, TGF-β/BMP signaling, and autophagy pathways. (**E**) Comparison of *C. burnetii* host kinase dependencies identified in this study with a previous kinase inhibitor screen (15), revealing shared regulators and high-confidence kinases uniquely identified through the current deconvolution approach.

Downstream process annotation and pathway enrichment analysis of these validated targets highlighted distinct functional configurations (**Figure 3D**). Supportive kinase networks were heavily dominated by proliferative and metabolic survival axes, including ERK/MAPK, RTK-survival, and AKT/PKC cascades, alongside structural cytoskeleton and vesicular trafficking machinery. In contrast, pathways mapped as suppressors of *C. burnetii* infection were tightly clustered around stress-activated and cell-autonomous innate immune responses, characterized by p38/JNK MAPKs, NF-κB/JAK/TLR signaling loops, and the TGF-β/Ephrin pathway. Intersecting our high-confidence hits with a foundational Genome-wide siRNA screen ^23^ demonstrated robust consensus overlap for historical core dependencies like EGFR, ERBB2, and MAPK14 (**Figure 3E**), while validating our platform’s unique capacity to isolate novel, cross-niche regulatory hubs like ALK2.

### *Coxiella burnetii* Intracellular Colonization Actively Drives and Relies Upon Host ALK2 Signaling

To isolate core signaling nodes required for *Coxiella* pathogenesis from cell-type-specific responses, we leveraged our comparative host cell framework. Across both host models, despite their inherent biological differences, we identified a robust, overlapping enrichment of the TGF-β/BMP superfamily. Based on this conserved activation, we focused on the functional role of ALK2 during vacuolar establishment. Longitudinal Western blot tracking of direct downstream effectors revealed that *C. burnetii* infection actively drives a progressive, step-wise activation of host ALK2 signaling, marked by a robust activation of p-SMAD1/5/9 levels during early stages of infection relative to uninfected controls (**Figure 4A**). In contrast, infection with a Dot/Icm type IV secretion system-deficient *C. burnetii* mutant failed to elicit significant p-SMAD1/5/9 phosphorylation throughout the infection time course, suggesting that activation of host ALK2 signaling is not a passive consequence of bacterial uptake but instead requires active manipulation of host signaling by Dot/Icm-translocated effectors (**Figure 4A)**.

**Figure 4.**
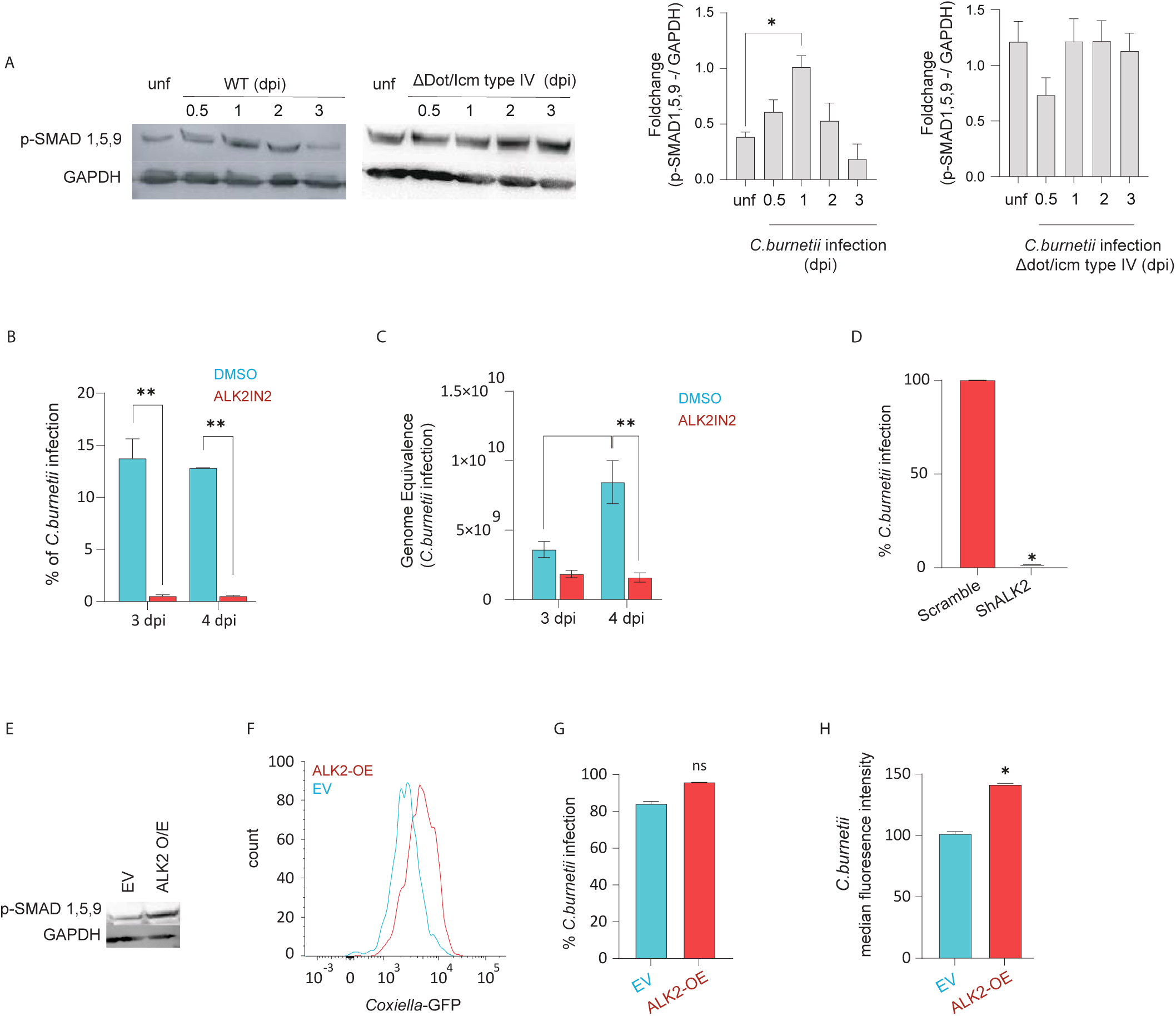
*C. burnetii* activates and depends on host ALK2 signaling for intracellular replication. (**A**) Immunoblot analysis and densitometric quantification of p-SMAD1/5/9 levels during infection with wild-type *C. burnetii* or a Dot/Icm type IV secretion-deficient mutant. Phosphorylation levels were normalized to GAPDH. (**B, C**) Pharmacological inhibition of ALK2 using ALK2IN2 reduces *C. burnetii* infection (**B**) and bacterial genome equivalents (**C**) at 3 and 4 days post-infection (dpi). (**D**) shRNA-mediated depletion of ALK2 impairs *C. burnetii* infection relative to scramble controls. (**E**) Immunoblot confirming increased p-SMAD1/5/9 levels following ALK2 overexpression (ALK2-OE). (**F**) Representative flow cytometry histograms of *C. burnetii*-GFP infection in empty vector (EV) and ALK2-overexpressing cells. (**G, H**) Quantification of infection frequency (G) and bacterial burden measured by GFP median fluorescence intensity (**H**) in EV and ALK2-OE cells. Data are representative of three independent biological experiments. Bars represent mean ± SD of technical replicates from the representative experiment shown. *P < 0.05, **P < 0.01; ns, not significant.

This functional dependency was independently confirmed using ALK2IN2, a highly selective small-molecule competitive inhibitor of the ALK2 kinase domain ^24^. ALK2IN2 treatment at a concentration that had no effect on cell viability (**Figure S3**) nearly abolished *C. burnetii* intracellular development at 3 and 4 dpi, completely blunting the log-linear expansion of absolute bacterial genome equivalence (**Figure 4B** and **4C**).

To orthogonally validate our pharmacological findings and confirm the functional requirement for host ALK2 during pathogenesis, we manipulated ALK2 expression using genetic silencing. Lentiviral shRNA-mediated knockdown of host ALK2 (shALK2) reduced the percentage of infection (*Coxiella-*positive cells) significantly compared to scramble controls (**Figure 4D**). Reciprocally, ectopic plasmid-driven overexpression of host ALK2 induced phosphorylation of SMAD (**Figure 4E**). While there is no significant alteration in the infection frequency, a significant increase in the *C. burnetti* burden (GFP fluorescence intensity) was observed, as evident from **Figure 4F, 4G** and **4H**. Together, these complementary pharmacological and genetic findings support our hypothesis, confirming that host ALK2 signaling is actively hijacked to drive successful vacuole establishment and ongoing CCV biogenesis.

### *Chlamydia* Opposes the ALK2 Signaling Axis to Support Inclusion Growth

Building on the enrichment of the TGF/SMAD pathway in our primary screen (**Figure S4**), we examined how the non-fusogenic, neutral-pH inclusion of *C. trachomatis* interacts with the host ALK2-SMAD pathway. In stark contrast to the progressive upregulation driven by *C. burnetii*, Western blot tracking of *C. trachomatis*-infected monolayers revealed a significant down-regulation of baseline host p-SMAD1/5/9 levels (**Figure 5A**), establishing that this pathogen actively blocks host ALK2 signaling outputs.

**Figure 5.**
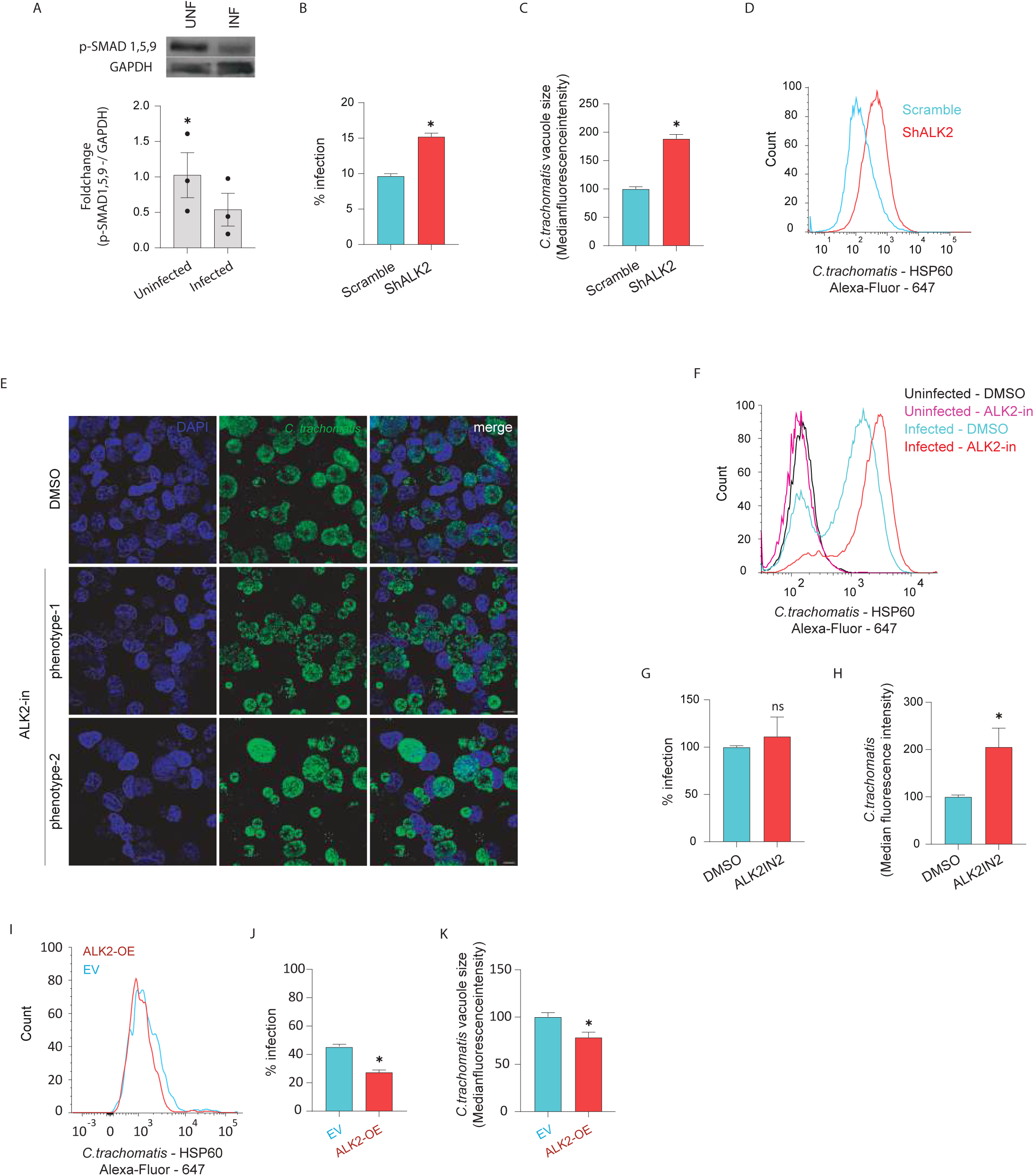
*C. trachomatis* suppresses host ALK2 signaling to promote inclusion development. (**A**) Immunoblot analysis and densitometric quantification of p-SMAD1/5/9 levels in uninfected and *C. trachomatis*-infected cells. Phosphorylation levels were normalized to GAPDH. (**B, C**) shRNA-mediated depletion of ALK2 increases *C. trachomatis* infection frequency (**B**) and inclusion size (**C**). (**D**) Representative flow cytometry histograms showing inclusion-associated HSP60 signal in scramble and shALK2 cells. (**E**) Representative confocal micrographs of infected cells treated with DMSO or ALK2IN2, showing distinct inclusion phenotypes following ALK2 inhibition. Scale bars, 10 μm. (**F**) Representative flow cytometry histograms of HSP60 staining in uninfected and infected cells treated with DMSO or ALK2IN2. (**G, H**) Quantification of infection frequency (**G**) and inclusion size (**H**) following pharmacological inhibition of ALK2. (I) Representative flow cytometry histograms of HSP60 signal in empty vector (EV) and ALK2-overexpressing (ALK2-OE) cells. (**J, K**) Quantification of infection frequency (**J**) and inclusion size (**K**) in EV and ALK2-OE cells. Data are representative of three independent biological experiments. Bars represent mean ± SD of technical replicates from the representative experiment shown. *P < 0.05; ns, not significant.

Consistent with this biochemical divergence, modulating host ALK2 yielded phenotypic outcomes that were the exact inverse of those observed during *C. burnetii* infection. Stable shRNA knockdown of host ALK2(shALK2) significantly enhanced *C. trachomatis* fitness, prompting a substantial increase in the overall percentage of infected cells and driving a dramatic expansion in the physical dimensions of the *Chlamydia* inclusion vacuole (**Figure 5B, 5C** and **5D**). This hyper-development phenotype was closely recapitulated by acute pharmacological inhibition using the selective ALK2 inhibitor, ALK2IN2, shifting host-pathogen dynamics in favor of the bacterium by augmenting infection rates and promoting inclusion expansion (**Figure 5E**). Immunofluorescence analysis following ALK2IN2 treatment revealed two distinct morphological phenotypes: host cells containing either a single hyper-enlarged inclusion or multiple smaller inclusions per cell. Flow cytometric analysis confirmed this heightened intracellular bacterial burden, demonstrating a significant increase in mean fluorescence intensity (MFI) among infected host cells (**Figures 5F-5H**).

Conversely, forcing pathway activation via ectopic receptor overexpression (ALK2-OE) severely restricted *C. trachomatis* pathogenesis significantly by reducing the percentage infection frequency and imposing a strict structural constraint that collapsed inclusion vacuole size (**Figure 5I, 5J** and **5K**). These results demonstrate a striking biological paradigm where distinct intracellular niches deploy opposing strategies to manipulate an identical host signaling node.

### Host ALK2 signaling mediates TFEB activation and LAMP1 recruitment to the *Coxiella*-containing vacuole

Having established that *C. burnetii* strictly depends on ALK2 signaling while *C. trachomatis* is antagonized by it, we sought to determine the downstream cellular mechanism driving this divergence. *C. burnetii* replication strictly requires the biogenesis and maturation of a highly acidic, phagolysosome-like vacuole (CCV). Because the transcription factor EB (TFEB) is a master host regulator of lysosomal biogenesis and autophagy, we hypothesized that ALK2 might regulate the pathogen’s niche via the TFEB axis.

To test this, we assessed the subcellular localization of TFEB in *C. burnetii*-infected cells following ALK2 blockade. In DMSO-treated control cells, infection robustly induced the nuclear translocation of TFEB. However, treatment with ALK2-IN-2 dramatically prevented TFEB nuclear translocation, retaining the transcription factor in the cytoplasm (**Figure 6A and 6C**). Because TFEB drives the expression of lysosomal proteins required for CCV maturation, we next examined the recruitment of the lysosomal-associated membrane protein 1 (LAMP1) to the bacterial vacuole. Immunofluorescence analysis revealed that while control CCVs were heavily decorated with LAMP1, pharmacological inhibition of ALK2 severely impaired LAMP1 recruitment to the *Coxiella* vacuoles (**Figure 6B and 6D**). Together, these data demonstrate that ALK2 signaling acts as a critical upstream checkpoint required for TFEB-mediated lysosomal biogenesis and subsequent CCV maturation, structurally linking the kinome screening data to the physical reality of the pathogen’s replicative niche.

**Figure 6.**
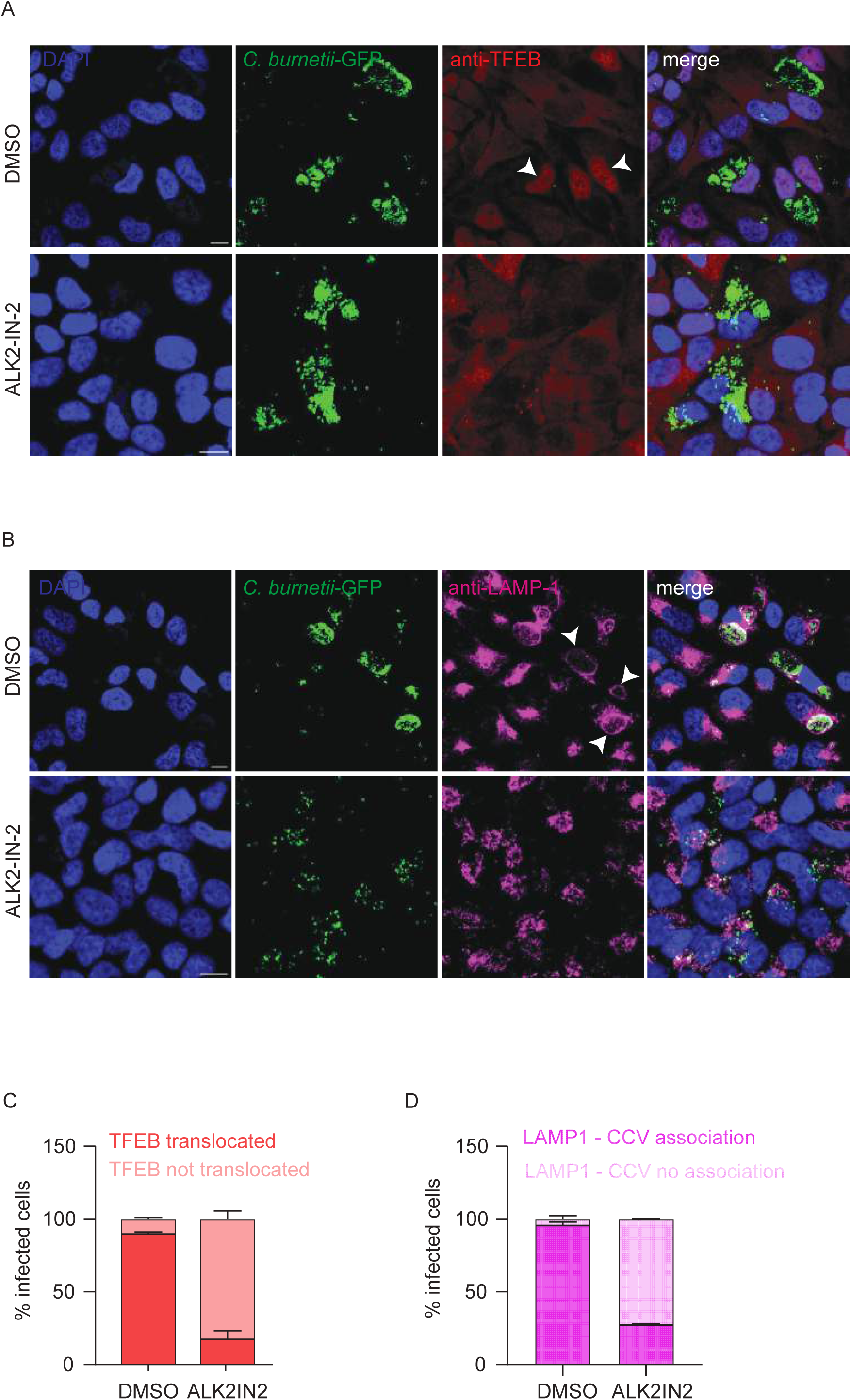
Host ALK2 signaling mediates TFEB nuclear translocation and LAMP1 recruitment to the *Coxiella*-containing vacuole. **(A)** Representative immunofluorescence images of *C. burnetii*-infected cells treated with DMSO (vehicle control) or the ALK2 inhibitor ALK2-IN-2. Cells were fixed at 72 hours post-infection (hpi) and stained for TFEB (red). Bacterial vacuoles are indicated by *C. burnetii*-GFP (green) and nuclei are stained with DAPI (blue). In DMSO-treated cells, TFEB robustly translocates to the nucleus, whereas ALK2-IN-2 treatment results in cytosolic retention of TFEB. Scale bar: 10 µm. Images are from a single representative experiment of three independent biological experiments. **(B)** Representative immunofluorescence images of *C. burnetii*-GFP (green) infected cells treated with DMSO or ALK2-IN-2 and stained for the late endosomal/lysosomal marker LAMP1 (red) at 72 hpi. Nuclear DNA is stained with DAPI (blue). Images are from a single representative experiment of three independent biological experiments. **(C)** Quantification of TFEB nuclear translocation from the experiments shown in (A). Data are represented as the percentage of infected cells displaying prominent nuclear versus cytosolic TFEB localization. **(D)** Quantification of LAMP1 association with the *Coxiella*-containing vacuole (CCV) from the experiments shown in (B). Data are represented as the percentage of infected cells containing vacuoles with distinct LAMP1 recruitment.

### Identification of host-targeted inhibitors of *C. burnetii* infection

In addition to identifying host kinases associated with *C. burnetii* infection, our machine learning framework was used to predict the efficacy of kinase inhibitors with previously characterized target selectivity profiles. Inhibitors were ranked according to their predicted effects on infection and vacuole development (**Figure 7A and B**). Among the compounds evaluated, Sunitinib was predicted to strongly impair infection in both HeLa (**Figure 7A and B**) and THP-1 (**Figure S5)**, whereas Jak inhibitor I was predicted to have limited activity. To test these predictions, infected cells were treated with each compound at a concentration that did not affect cell viability (**Figure S6**), and infection phenotypes were quantified. Consistent with model predictions, Sunitinib significantly reduced both the proportion of infected cells and CCV size relative to vehicle-treated controls, whereas Jak inhibitor I produced little effect (**Figure 7C, D** and **E**). Taken together, we were able to predict, and test host kinases involved in infection and also identify novel host-targeted inhibitors that are effective at dramatically reducing *C. burnetii* infection.

**Figure 7.**
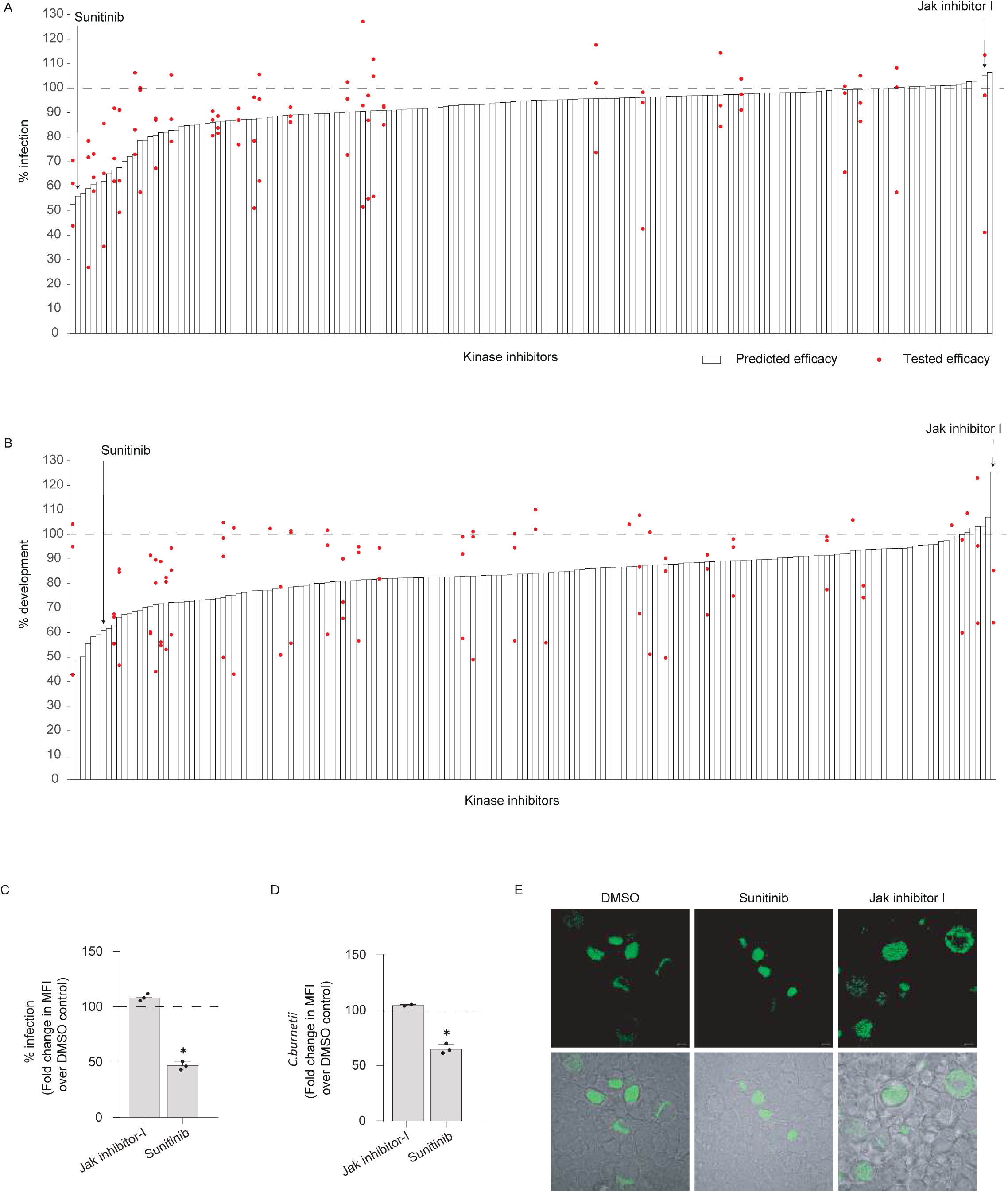
Machine learning-guided prediction identifies host-targeted inhibitors of *C. burnetii* infection. (**A** and **B**) Comparison of predicted and experimentally measured effects of kinase inhibitors on *C. burnetii* infection (**A**) and intracellular development (**B**). Predicted efficacies were generated using elastic net regression models trained on kinase inhibitor activity profiles and infection phenotypes, while red dots indicate experimentally measured values. Sunitinib and Jak inhibitor I are highlighted as representative compounds predicted to decrease or minimally affect infection, respectively. (**C** and **D**) Experimental validation of model predictions showing the effects of Sunitinib and Jak inhibitor I on infection frequency (**C**) and Coxiella-containing vacuole (CCV) size (**D**), expressed relative to DMSO-treated controls. (**E**) Representative fluorescence and phase-contrast images of *C. burnetii*-GFP-infected cells treated with DMSO, Sunitinib, or Jak inhibitor I. Scale bars, 10 μm. Data are representative of three independent biological experiments. Bars represent mean ± SD of technical replicates from the representative experiment shown. *P < 0.05.

## Discussion

Intracellular pathogens depend on host cell pathways to acquire nutrients, maintain their replicative niches, and avoid host defense mechanisms. A long-standing question in the field is whether different intracellular pathogens exploit a common set of host regulatory networks or rely on pathogen-specific strategies. In this study, we combined a multi-pathogen kinase inhibitor screen with machine learning-based polypharmacology deconvolution to systematically identify host pathways that regulate infection by *C. burnetii, C. trachomatis,* and *S. enterica*. This approach allowed us to move beyond nominal drug targets and generate a comparative map of host kinase dependencies across distinct intracellular pathogens.

Our analyses revealed substantial differences in host kinase dependence among the three pathogens. *C. burnetii* was highly sensitive to host kinase perturbation, whereas *C. trachomatis* and *S. enterica* showed more restricted responses. These differences likely reflect the distinct biological demands of their intracellular niches. The Coxiella-containing vacuole is a highly fusogenic lysosome-derived compartment that undergoes continuous fusion with the endolysosomal system and requires sustained membrane trafficking and metabolic support for expansion. Moreover, *C. burnetii* completes a prolonged intracellular developmental cycle over approximately 5 days, during which it remains dependent on host pathways for vacuole maintenance and bacterial replication ^5^. In contrast, *C. trachomatis* and *S. enterica* complete much shorter intracellular replication cycles, typically within 24-48 hours, and their vacuoles interact with a more restricted subset of host trafficking pathways. Consequently, perturbation of host kinase signaling is expected to exert a greater cumulative effect on *C. burnetii* than on the other pathogens.

Polypharmacology deconvolution using elastic net regression enabled the reconstruction of host signaling networks that regulate *C. burnetii* infection in both epithelial cells and macrophages. Pathways associated with infection included ERK/MAPK, receptor tyrosine kinase, and AKT/PKC signaling, whereas stress-responsive pathways such as p38/JNK and NF-kB signaling were associated with restriction of infection. The strong overlap between our predictions and previously identified host regulators ^14^ supports the robustness of this approach and demonstrates the value of kinase inhibitor profiling for identifying host pathways that contribute to intracellular infection.

A major finding of this study is the identification of ALK2 signaling as a host pathway that is regulated in opposite directions by different intracellular pathogens. ALK2 (Activin A receptor, type I), canonically known as a bone morphogenetic protein (BMP) type I receptor critical for skeletal development and tissue homeostasis ^32^, emerges here as an unexpected central fulcrum of intracellular pathogenesis. During *C. burnetii* infection, ALK2 signaling was progressively activated, resulting in increased phosphorylation of SMAD1/5/9. Genetic silencing, chemical inhibition, or dominant suppression of ALK2 impaired bacterial replication and restricted CCV expansion, whereas ALK2 overexpression enhanced vacuole development. These findings indicate that *C. burnetii* actively exploits host ALK2 signaling to support intracellular growth. In contrast, *C. trachomatis* suppressed basal ALK2 signaling, and inhibition of ALK2 enhanced infection, whereas pathway activation restricted inclusion development. These observations suggest that a single host signaling pathway can have opposing effects depending on the biological requirements of the pathogen. More broadly, they highlight how intracellular pathogens can reshape the same host regulatory network in different ways to optimize their intracellular niche.

A major conceptual advance of this study is the delineation of why ALK2 signaling yields divergent outcomes for different intracellular pathogens. Our mechanistic data reveal that ALK2 signaling functions upstream of TFEB nuclear translocation and subsequent LAMP1 recruitment to the pathogen vacuole (**Figure 6**). This perfectly aligns with the distinct evolutionary strategies of the pathogens screened. *C. burnetii* obligatorily requires a heavily acidified, LAMP1-positive phagolysosomal niche for its metabolism and replication. By co-opting the ALK2-SMAD axis, *C. burnetii* ensures the TFEB-driven lysosomal biogenesis ^33^ required to sustain its massive vacuole. In stark contrast, *C. trachomatis* resides in a neutral, Golgi-derived inclusion and actively secretes effector proteins to *prevent* fusion with lysosomes^6^. Therefore, an active ALK2-TFEB axis which promotes lysosomal abundance and trafficking would be fundamentally hostile to the *Chlamydia* inclusion. This mechanistic dichotomy beautifully illustrates how overlapping host signaling hubs are differentially exploited or avoided depending on the specific structural and biochemical requirements of the pathogen’s microenvironment.

Finally, our study demonstrates the utility of machine learning-guided host-directed drug discovery. By integrating kinase selectivity profiles with experimentally derived infection phenotypes, the model successfully identified Sunitinib as a compound with anti-infective activity against *C. burnetii* (**Figure 7**). Experimental validation confirmed that Sunitinib reduced both infection burden and CCV development without detectable host toxicity. More broadly, these results demonstrate that polypharmacology-based models can be used to prioritize host-directed therapeutics and identify candidate compounds for repurposing. Because host pathways evolve more slowly than pathogen targets, such approaches may provide a complementary strategy for developing treatments against intracellular infections that are less susceptible to the emergence of pathogen resistance.

## Resource Availability

### Lead contact

Further information and requests for resources and reagents should be directed to and will be fulfilled by the lead contact, Kamalakannan Vijayan

### Materials availability

All unique reagents generated in this study are available from the lead contact with a completed Materials Transfer Agreement.

### Data and Code availability

All data reported in this paper will be shared by the lead contact upon request. Any additional information required to reanalyze the data reported in this paper is available from the lead contact upon request. Custom scripts used for kinase regression analysis and downstream computational analyses are provided in the Supplementary Information.

### Use of Artificial Intelligence Tools

ChatGPT (OpenAI) was used during manuscript preparation to assist with language editing and improve readability. All scientific content, analyses, interpretations, and conclusions were verified by the authors, who take full responsibility for the final manuscript.

## Experimental Model and Subject Details

### Cell culture

HeLa cells were maintained in Dulbecco’s Modified Eagle Medium (DMEM; Gibco, 10569-010) supplemented with 10% fetal bovine serum (FBS) (Cytiva:SV30160.03) and penicillin-streptomycin (Cytiva:SV30010) at 37°C in a humidified atmosphere containing 5% CO₂. Cells were routinely passaged at 70–80% confluency. THP-1 cells were maintained in RPMI 1640 medium (Gibco, A10491-01) supplemented with 10% FBS and penicillin-streptomycin under identical culture conditions. Differentiation into macrophage-like cells was induced by treatment with 200 nM 12-O-tetradecanoylphorbol-13-acetate (TPA; Cell Signaling Technology, 4174S).

### Pathogen propagation and infection

#### Coxiella burnetii

The pGFP-expressing and *icmL*::Tn strains of *Coxiella burnetii* were propagated in Acidified Citrate Cysteine Medium-2 (ACCM-2; Sunrise Science, 4700-003) supplemented with kanamycin (375 μg mL⁻¹). Cultures were maintained at 37°C in a modular hypoxia chamber containing 5% CO₂, 2.5% O₂, and 92.5% N₂. After 6 days of growth, bacteria were harvested by centrifugation and resuspended in DMEM containing 5% FBS prior to infection.

#### Chlamydia trachomatis

*Chlamydia trachomatis* serovar L2 was propagated in HeLa cells at a multiplicity of infection (MOI) of 1. Infected cells were mechanically disrupted, and bacterial elementary bodies were purified by sequential low-speed and high-speed centrifugation. Purified bacteria were resuspended in SPG buffer (7.5% sucrose, 0.052% KH₂PO₄, 0.122% Na₂HPO₄, and 0.072% L-glutamate), aliquoted, and stored at −80°C. Infectious titres were determined before use, and infections were performed at the indicated MOI.

#### Salmonella enterica serovar Typhimurium

*Salmonella enterica serovar Typhimurium* strain SL1344 expressing mCherry was recovered from glycerol stocks and cultured in Luria–Bertani (LB) (Himedia; M1245-500G) medium. Exponential-phase cultures (OD₆₀₀ = 0.9) were used to infect HeLa cells at an MOI of 100. One hour after infection, extracellular bacteria were removed by washing with phosphate-buffered saline (PBS) (Cytiva: SH30378.02), and cells were maintained in culture medium containing 100 µg mL⁻¹ gentamicin (Himedia: A005-20ML).

## Method Details

### Kinase inhibitor screening

For *C. burnetii* infection assays, HeLa cells were seeded at 1 × 10⁵ cells per well in 24-well plates, whereas TPA-differentiated THP-1 macrophages were seeded at 7.5 × 10⁴ cells per well in 96-well plates. Cells were infected with *C. burnetii* at an MOI of 100 and treated with kinase inhibitors (0.5 μM) at 6 h post-infection. Fresh inhibitor was added at 2 and 4 days post-infection. For *C. trachomatis* assays, HeLa cells were infected at an MOI of 1 and treated with kinase inhibitors (0.5 μM) at 4 h post-infection. Cells were fixed with 4% paraformaldehyde (Himedia: TCL119-100ML) prior to analysis.

### Quantification of intracellular infection

Intracellular pathogen burden was quantified using pathogen-specific fluorescence-based assays. Infection by pGFP-expressing C. burnetii in HeLa cells was quantified by flow cytometry, whereas infection in THP-1 macrophages was quantified using fluorescence plate-reader measurements. For *C. trachomatis* infection, cells were fixed, permeabilized with 0.1% Triton X-100 (Sigma T8787), and blocked in 5% bovine serum albumin (BSA) in PBS (Himedia-ML184-500ML). Chlamydial inclusions were detected using an anti-HSP60 monoclonal antibody followed by an Alexa Fluor 488-conjugated secondary antibody (Invitrogen: A10680). Infection levels were quantified by fluorescence microscopy. Intracellular mCherry-*Salmonella* infection was quantified by flow cytometry.

### shRNA-mediated gene silencing

Lentiviral particles were generated in HEK293T cells by co-transfection of pLKO.1 shRNA constructs together with psPAX2 and pMD2.G packaging plasmids using Lipofectamine™ 3000 Transfection Reagent (Invitrogen: L3000150). Viral supernatants were collected 48 h after transfection and stored at −80°C.

HeLa cells were transduced in the presence of polybrene (1 μg mL⁻¹) (Sigma: TR-1003-G) and selected with puromycin (2 μg mL⁻¹) (Gibco: A1113803) beginning 24 h after transduction.

Selection was maintained for 5–6 days before downstream analyses. Knockdown efficiency was confirmed by western blotting.

### Western blotting

Cells were lysed in RIPA buffer (Sigma:20-188) supplemented with protease inhibitors, and protein concentrations were determined using a bicinchoninic acid (BCA) (Invitrogen: A55860) assay. Equal amounts of protein (20–30 μg) were separated by SDS–PAGE and transferred to polyvinylidene difluoride membranes.

Membranes were blocked in 5% BSA prepared in Tris-buffered saline containing 0.1% Tween-20 (TBST) and incubated overnight at 4°C with primary antibodies. Following washing, membranes were incubated with horseradish peroxidase-conjugated secondary antibody (Invitrogen: 31460), for 1 h at room temperature. Signal detection was performed using enhanced chemiluminescence reagents (Miilipore: WBLUC0100, WBLUF0100). Band intensities were quantified using ImageJ and normalized to GAPDH.

### Immunofluorescence microscopy

Cells were cultured on L-polylysine-coated coverslips and infected with the indicated pathogens. For *C. trachomatis* infection, cells were fixed, permeabilized with 0.1% Triton X-100, blocked with 5% BSA, and stained with an anti-HSP60 antibody (Invitrogen: MA3-023) followed by an Alexa Fluor 488-conjugated secondary antibody (Invitrogen: A10680). Nuclei were counterstained with DAPI (1 μg mL⁻¹) (Invitrogen: D1306). For pGFP-expressing *C. burnetii* infection, cells were fixed, permeabilized (0.1% Triton-X), and counterstained with DAPI. Coverslips were mounted using Fluoromount (Invitrogen:00-4958-02) and imaged using an Olympus FV3000 confocal microscope.

### Genome equivalence assay

HeLa cells were infected with pGFP-expressing *Coxiella burnetii* at an MOI of 100. At 6 h post-infection, cells were treated with ALK2IN2 (HY-112815), whereas vehicle control wells received an equivalent volume of DMSO (D4540-100ML). Fresh inhibitor was added every 48 h throughout the experiment. At the indicated time points, infected cells were harvested and divided into two fractions. For genomic quantification, total DNA was isolated using the Origin Genomic DNA Isolation Kit (ODP301-03), and *C. burnetii* genome equivalents were quantified by quantitative PCR (qPCR) using *dotA*-specific primers. The second fraction was fixed with 4% paraformaldehyde and analyzed by flow cytometry to determine infection levels.

### Cell viability assay

Cell viability was assessed using the MTT reduction assay. Cells were seeded in 96-well plates at a density of 1 × 10⁴ cells per well and treated with kinase inhibitors for 72 h. Fresh medium and inhibitors were replenished every 24 h. Staurosporine (62996-74-1) (1 μM) or 5% DMSO served as positive controls for cytotoxicity. At the end of the treatment period, cells were incubated with MTT reagent (5 mg mL⁻¹ stock solution) for 3.5 h at 37°C. Formazan crystals were solubilized in DMSO, and absorbance was measured at 540 nm using a Tecan multimode microplate reader. Cell viability was calculated relative to vehicle-treated controls.

### Kinase regression analysis

Kinase regression (KiR) analysis was performed as previously described by Glennon et al.^26^. Briefly, intracellular infection measurements, including percent infection and median fluorescence intensity (MFI), were normalized to DMSO-treated controls and integrated with biochemical profiling data describing the activity of 38 kinase inhibitors against 291 recombinant protein kinases. Elastic net regression was implemented using the glmnet package in R with an elastic net mixing parameter (α) of 0.8. Regression coefficients were used to identify host kinases associated with intracellular infection and to predict the phenotypic effects of untested kinase inhibitors.

### Statistical analysis

Statistical analyses were performed using GraphPad Prism version 10. Unless otherwise indicated, comparisons between two groups were performed using two-tailed Welch’s t-tests. Statistical significance was defined as P < 0.05. The number of biological replicates, technical replicates, sample sizes, and statistical tests used for individual experiments are provided in the corresponding figure legends.

## Acknowledgements

The kinase inhibitor library was kindly gifted by Dr Alexis Kaushansky, Seattle Children’s Research Institute, Seattle, Washington, USA. This work was supported by the Science and Engineering Research Board-Anusandhan National Research Foundation (SERB-ANRF) (SRG/2023/001874) to K.V., the Department of Health Research-Indian Council of Medical Research (DHR-ICMR) (2022-1021) to K.V., the DBT-Wellcome Trust India Alliance Intermediate Fellowship (IA/I/23/2/506998) to K.V., and intramural funding from the Indian Institute of Science Education and Research Thiruvananthapuram (IISER Thiruvananthapuram) to K.V. *Chlamydia trachomatis serovar L2 strain* was a kind gift from Karthika Rajeeve, Rajiv Gandhi Center for Biotechnology, Thiruvananthapuram, Kerala, India. We thank Vaisak Mohan and Sandra S N for their help in executing the codes used in this study. We thank Anjana Narayanan for her assistance in carrying out the screen.

---

**Supplementary Figure 1.**
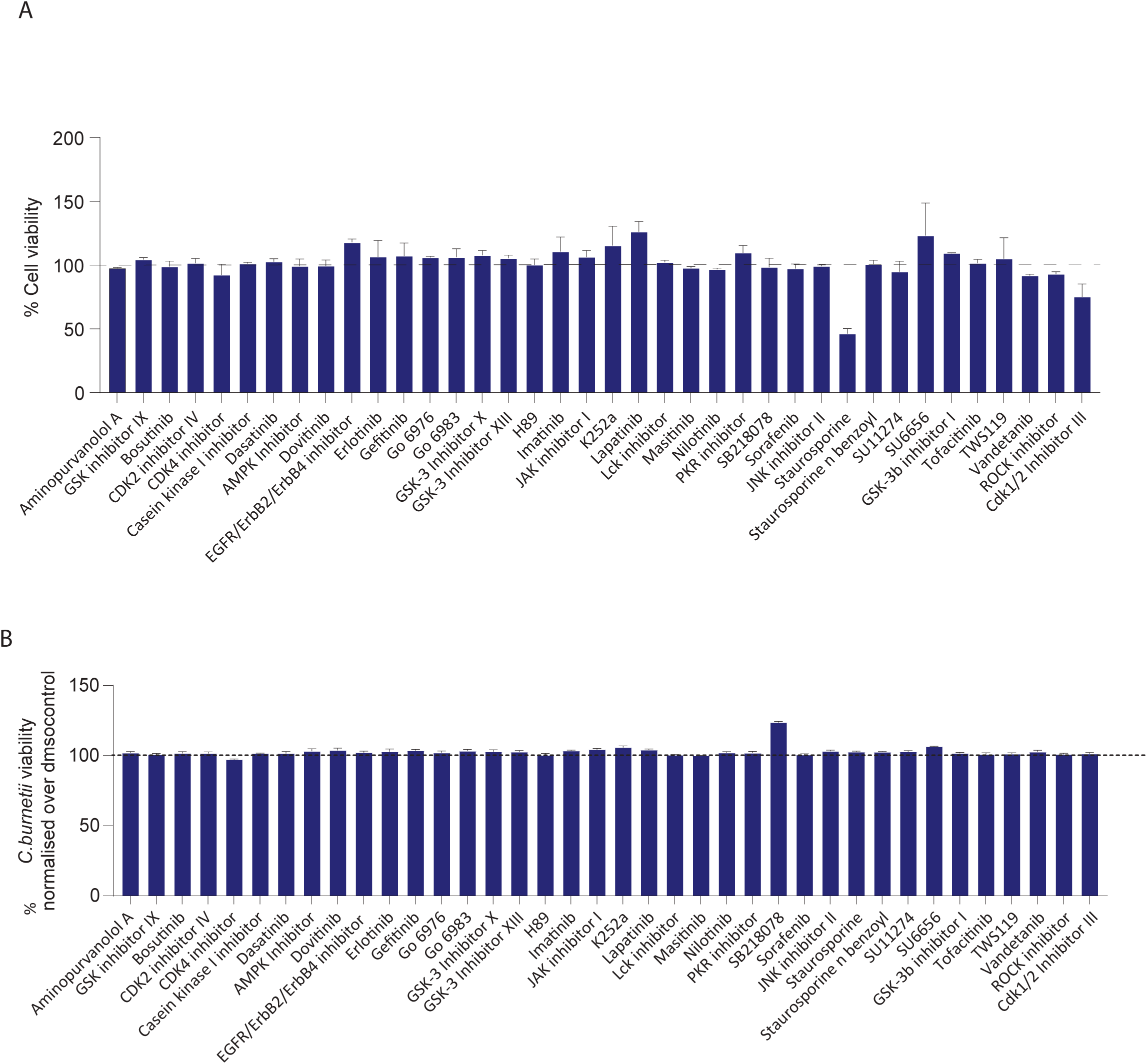
(A) Cytotoxicity assessment of the kinase inhibitor library by MTT assay. Host cell viability was assessed by MTT assay following treatment with the indicated kinase inhibitors at the concentrations used in the primary chemical screen. Cell viability is expressed as a percentage relative to DMSO-treated control cells, which were normalized to 100% (dashed line). Most inhibitors did not significantly affect host cell viability under the screening conditions, indicating that their effects on infection were not attributable to nonspecific cytotoxicity. Data are representative of three independent biological experiments. Bars represent mean ± SD of technical replicates from the representative experiment shown. *P < 0.05.

**Supplementary Figure 2.**
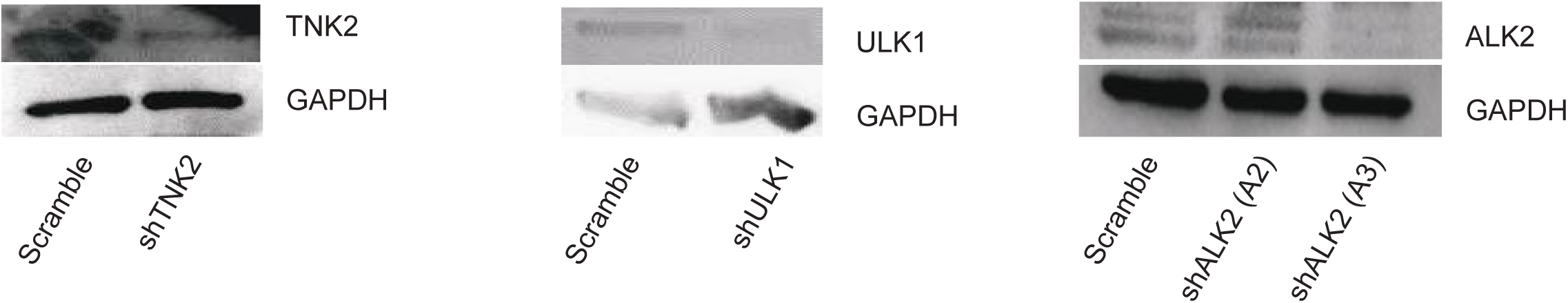
Validation of stable lentiviral shRNA-mediated knockdown of candidate host kinases. Representative immunoblots confirming stable lentiviral shRNA-mediated depletion of TNK2, ULK1, and ALK2 in host cells. GAPDH was used as the loading control. Two independent shRNAs targeting ALK2 (A2 and A3) were used to validate efficient ALK2 depletion. Representative immunoblots from three independent biological experiments are shown.

**Supplementary Figure 3.**
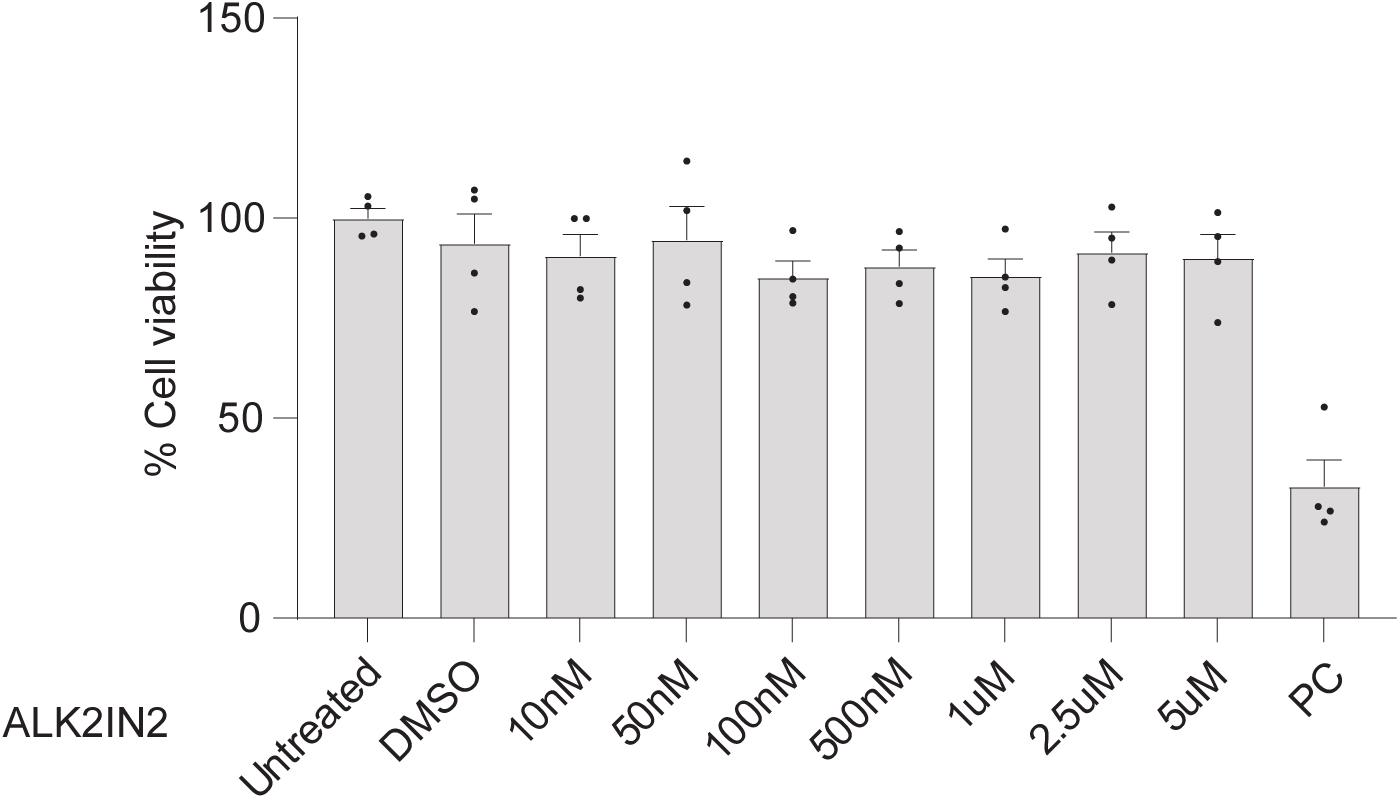
Cytotoxicity assessment of the ALK2 inhibitor by MTT assay. Host cell viability was determined by MTT assay following treatment with increasing concentrations of the Alk2-IN-2 (10 nM-5 μM). Cell viability is expressed as a percentage relative to untreated control cells. The inhibitor exhibited minimal cytotoxicity across the concentration range tested, whereas the positive control (PC) markedly reduced cell viability. Data are presented as mean ± SD from three independent biological experiments (n = 3).

**Supplementary Figure 4.**
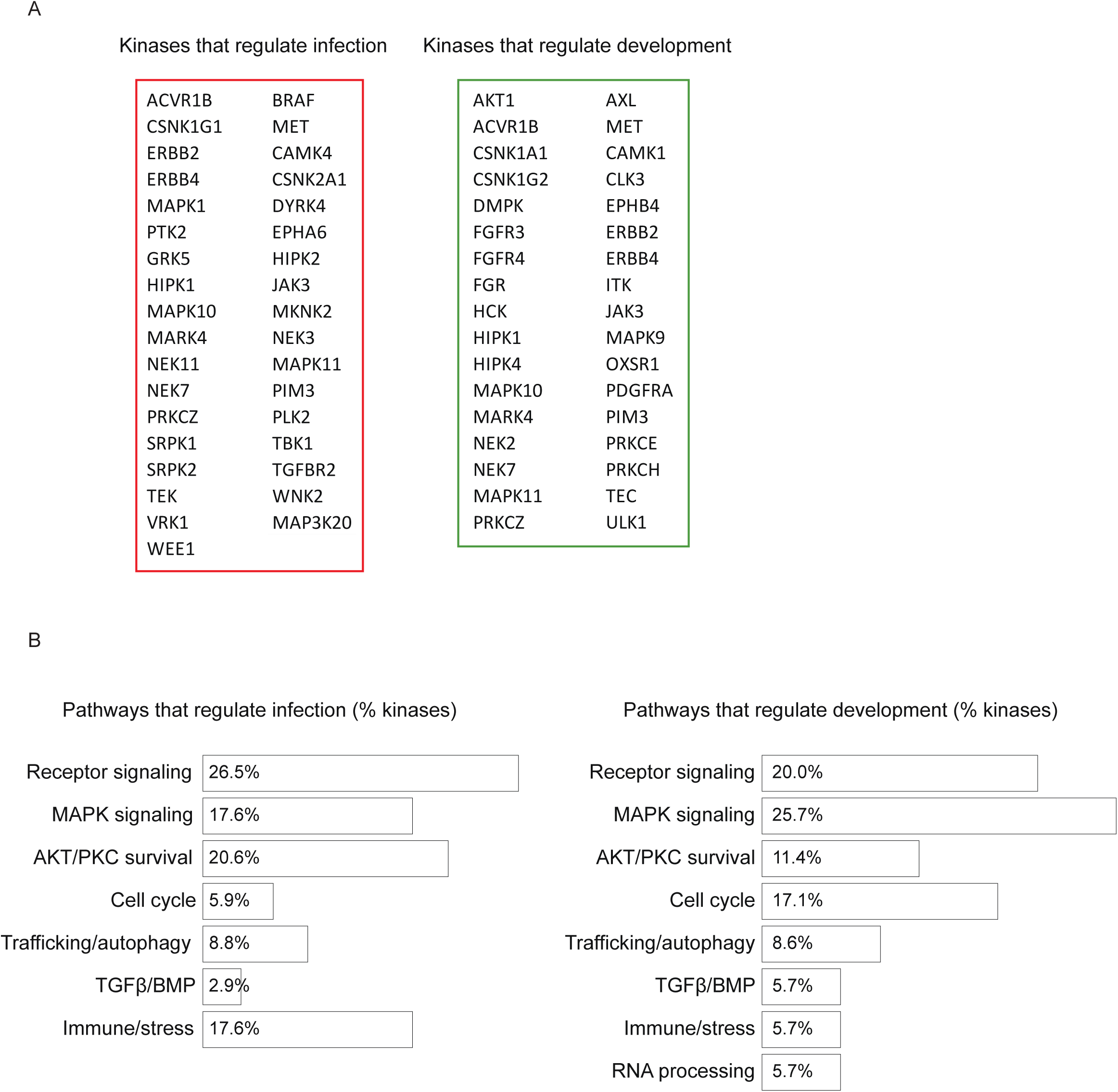
Functional classification of host kinases regulating *Chlamydia* infection and intracellular development. **(A)** Kinases identified from the primary kinase inhibitor screen that regulate infection frequency (left) or intracellular development (right). **(B)** Functional categorization of the identified kinases into major signaling pathways, including receptor signaling, MAPK signaling, AKT/PKC survival signaling, cell cycle regulation, trafficking/autophagy, TGFβ/BMP signaling, immune/stress signaling, and RNA processing. Percentages indicate the proportion of kinases assigned to each functional category within the infection- or development-associated kinase sets.

**Supplementary Figure 5.**
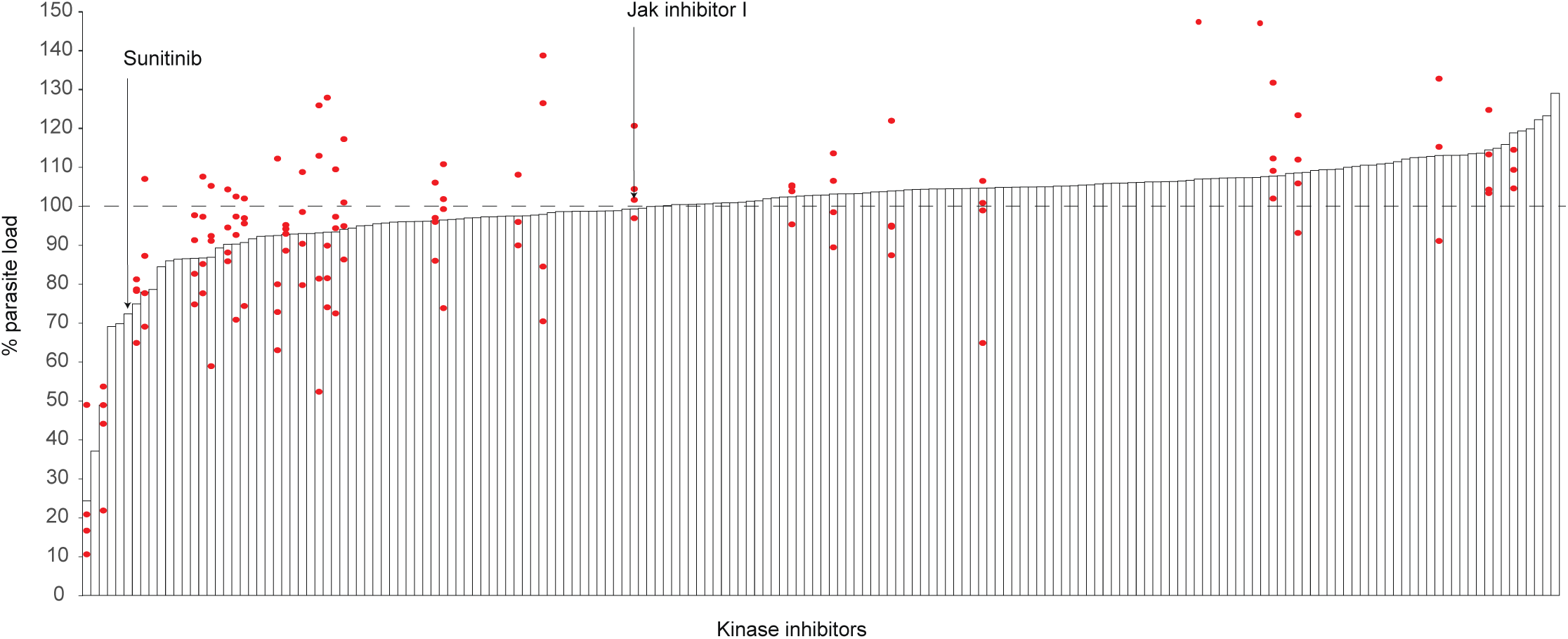
Kinase inhibitor screen identifying host kinases regulating *C. burnetii* infection in THP-1 cells. Bacterial load following treatment with the 178-compound kinase inhibitor library in THP-1 cells. Bacterial load is expressed as a percentage relative to DMSO-treated control cells (dashed line, 100%). Compounds are ranked according to their effect on infection. Inhibitors that reduce load are positioned to the left of the dashed line, whereas compounds that enhance infection are positioned to the right. Sunitinib and JAK Inhibitor I, selected for subsequent validation, are indicated.

**Supplementary Figure 6.**
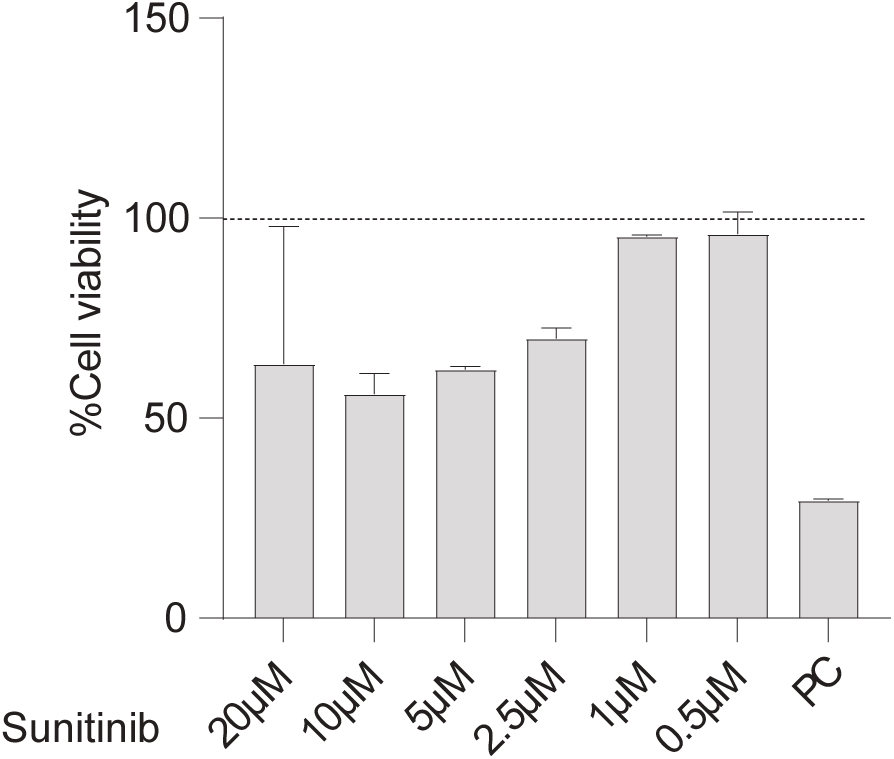
Cytotoxicity assessment of Sunitinib by MTT assay. Host cell viability was determined by MTT assay following treatment with increasing concentrations of Sunitinib (0.5–20 μM). Cell viability is expressed as a percentage relative to untreated control cells. Sunitinib exhibited minimal cytotoxicity at concentrations up to 20 μM under the experimental conditions, whereas the positive control (PC) markedly reduced cell viability. Data are presented as mean ± SD from three independent biological experiments (n = 3).

## Supplementary Note 1

Below is the script used to perform the regression analysis included in the manuscript.

The script can be run in R program

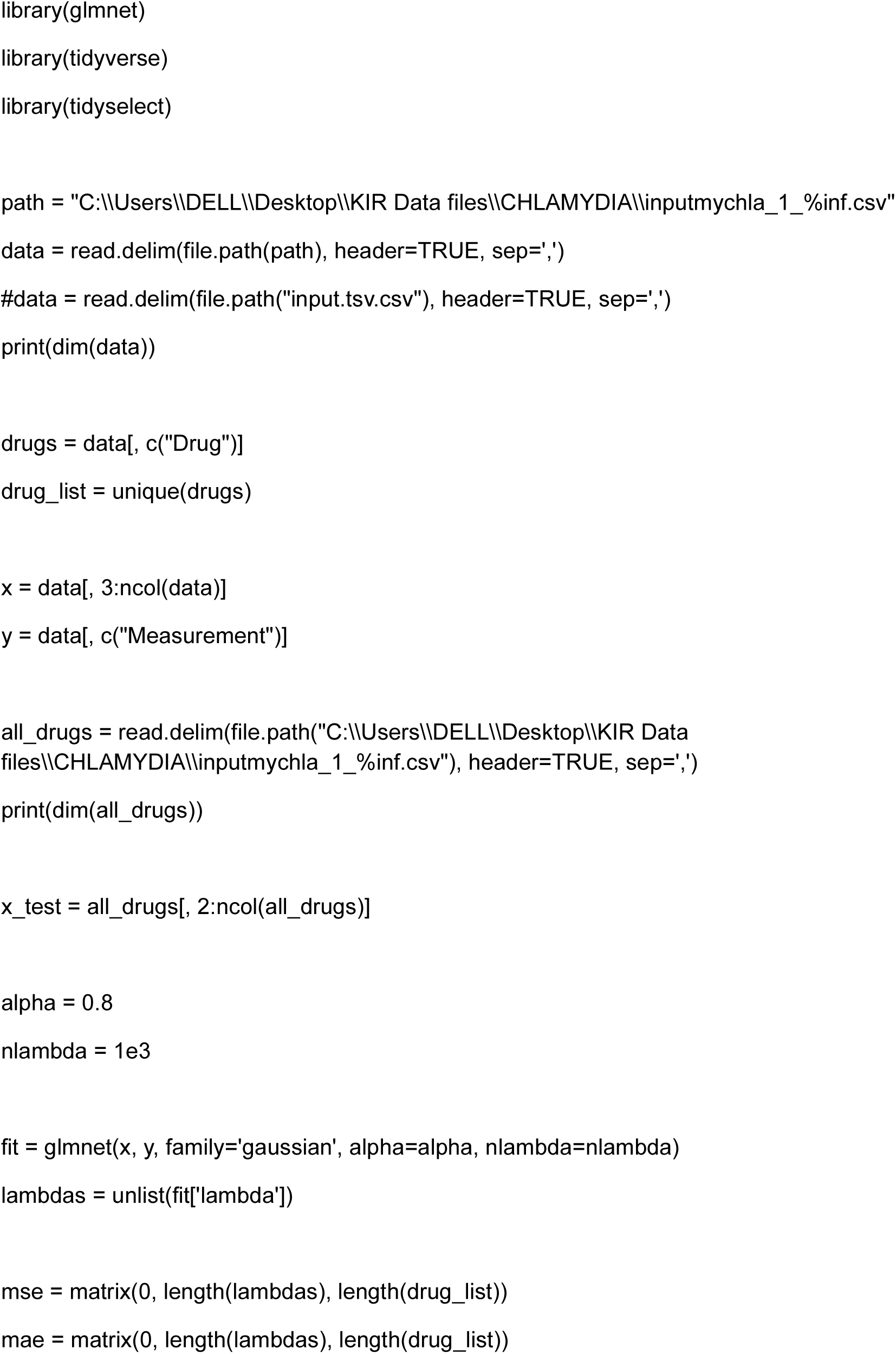
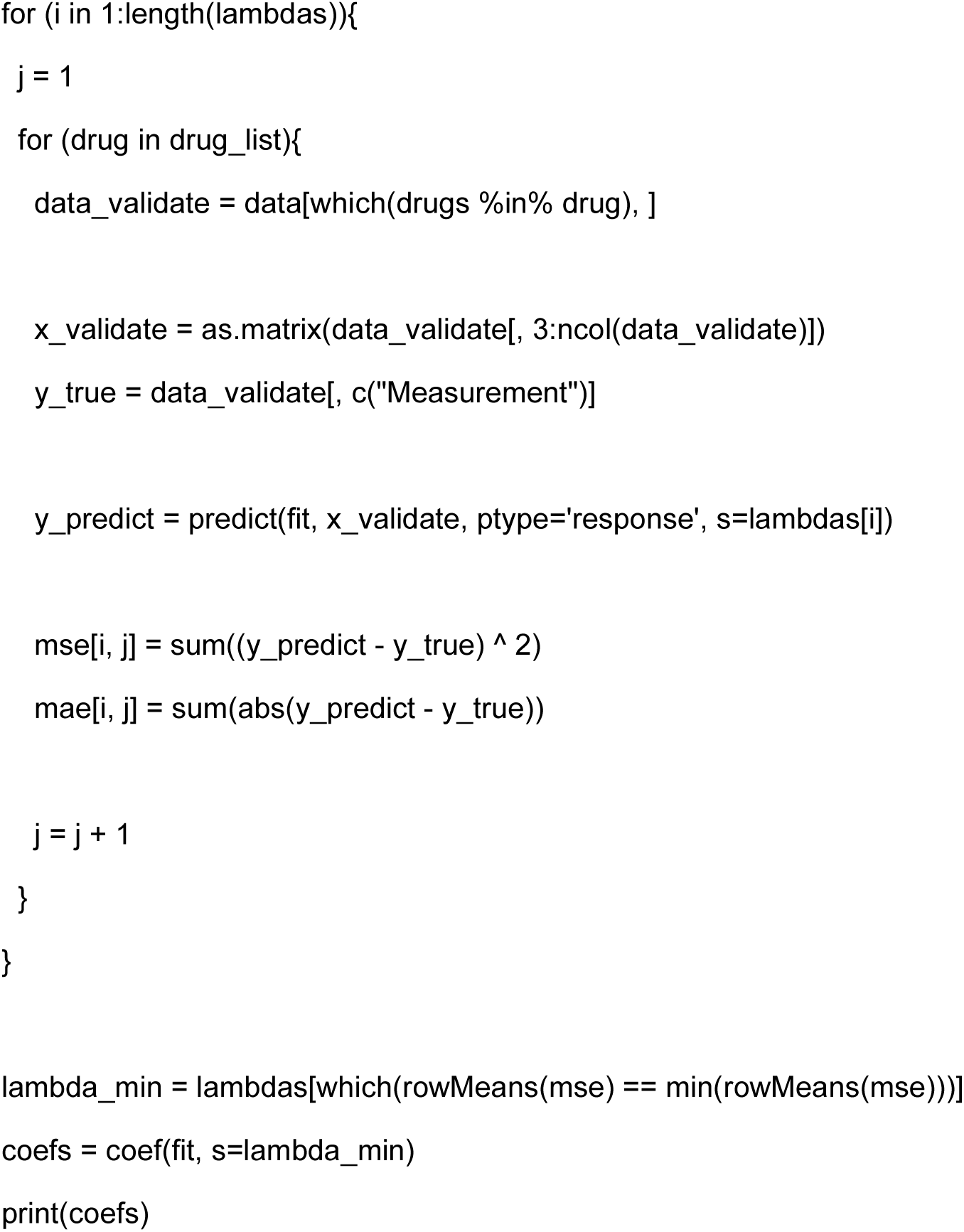

## Supplementary Note 2

Below is the script used to predict the effect of 178 inhibitors on infection frequency and development mentioned in the manuscript. Related to Figure 6.

The script can be run in Python program

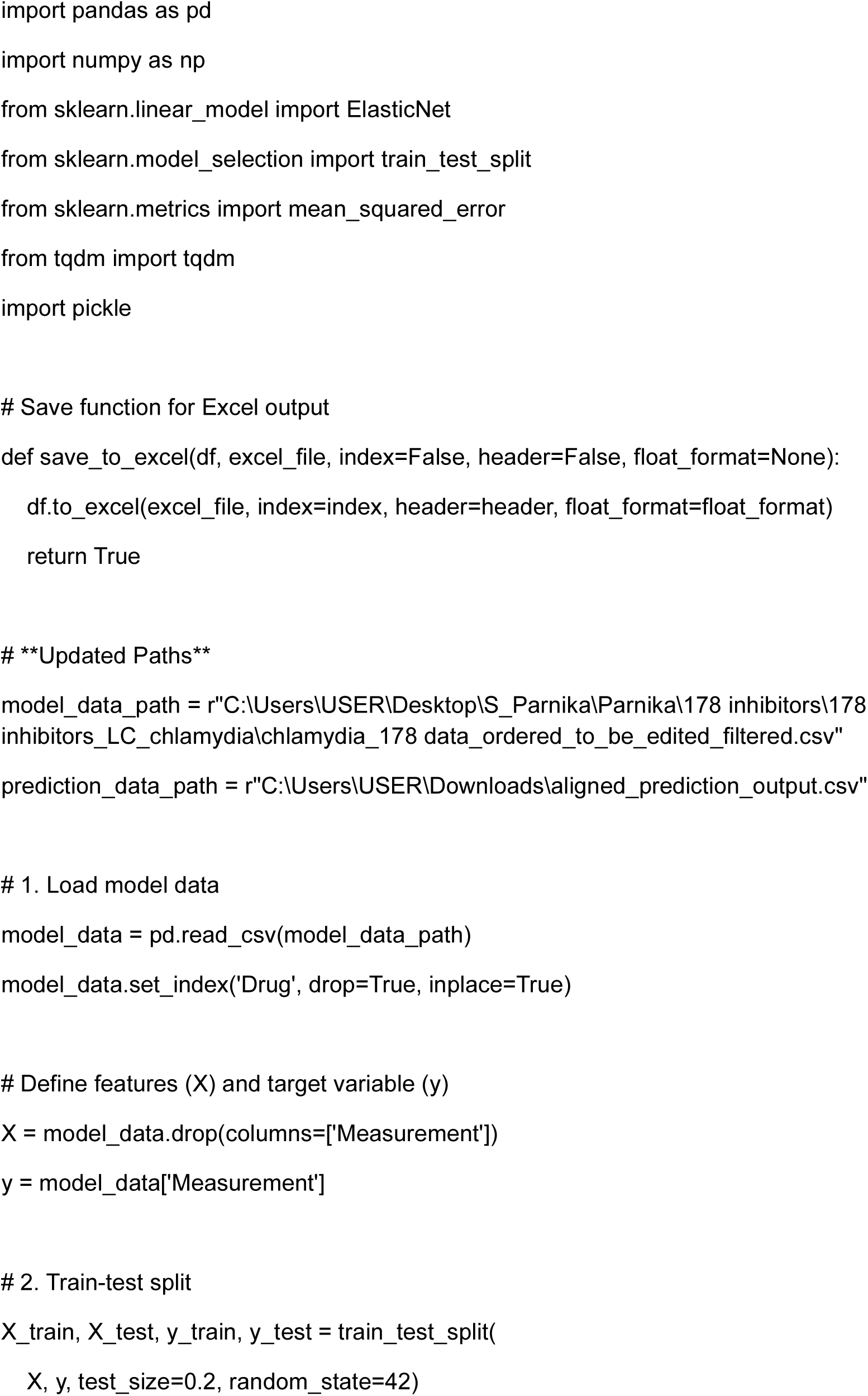
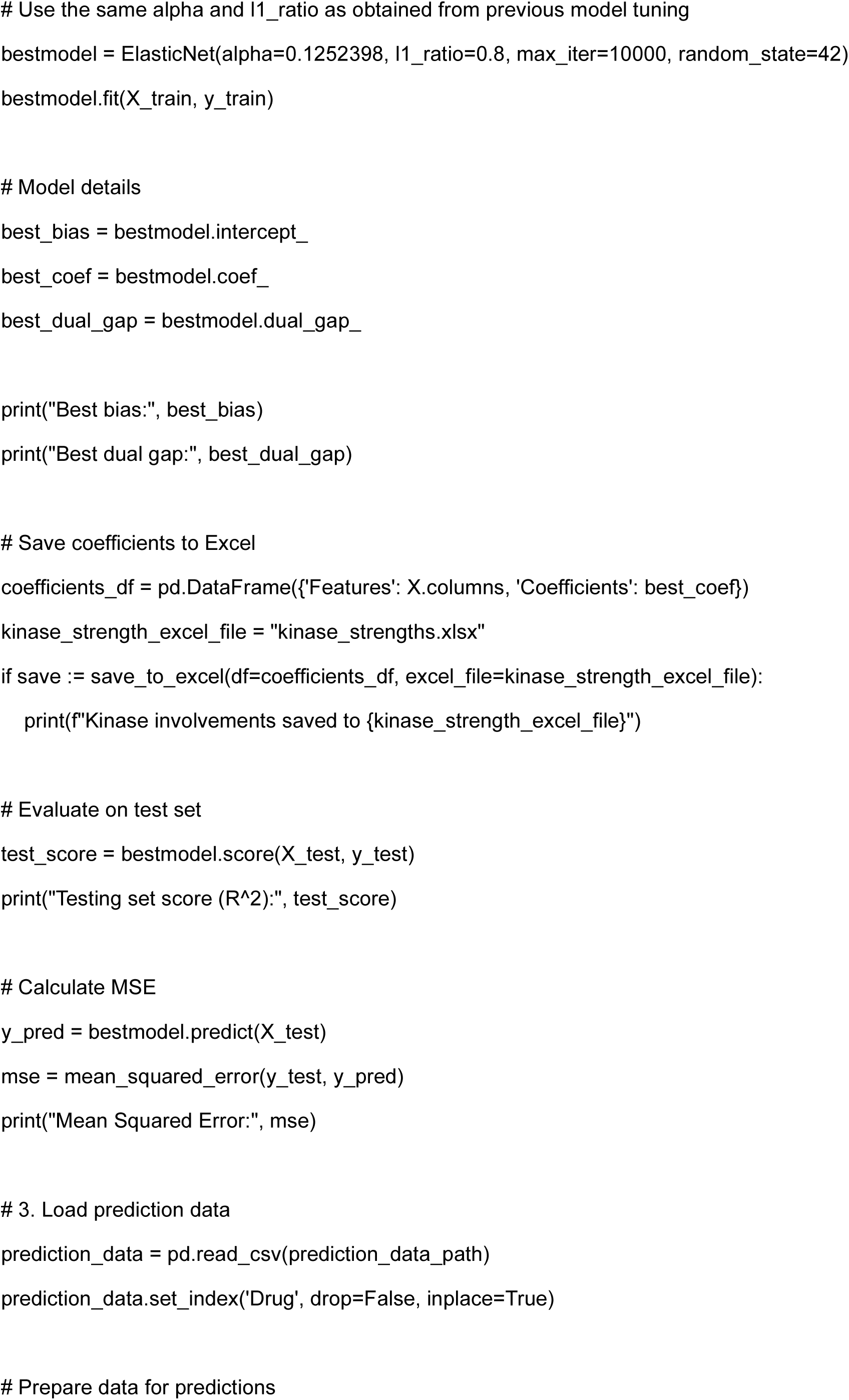
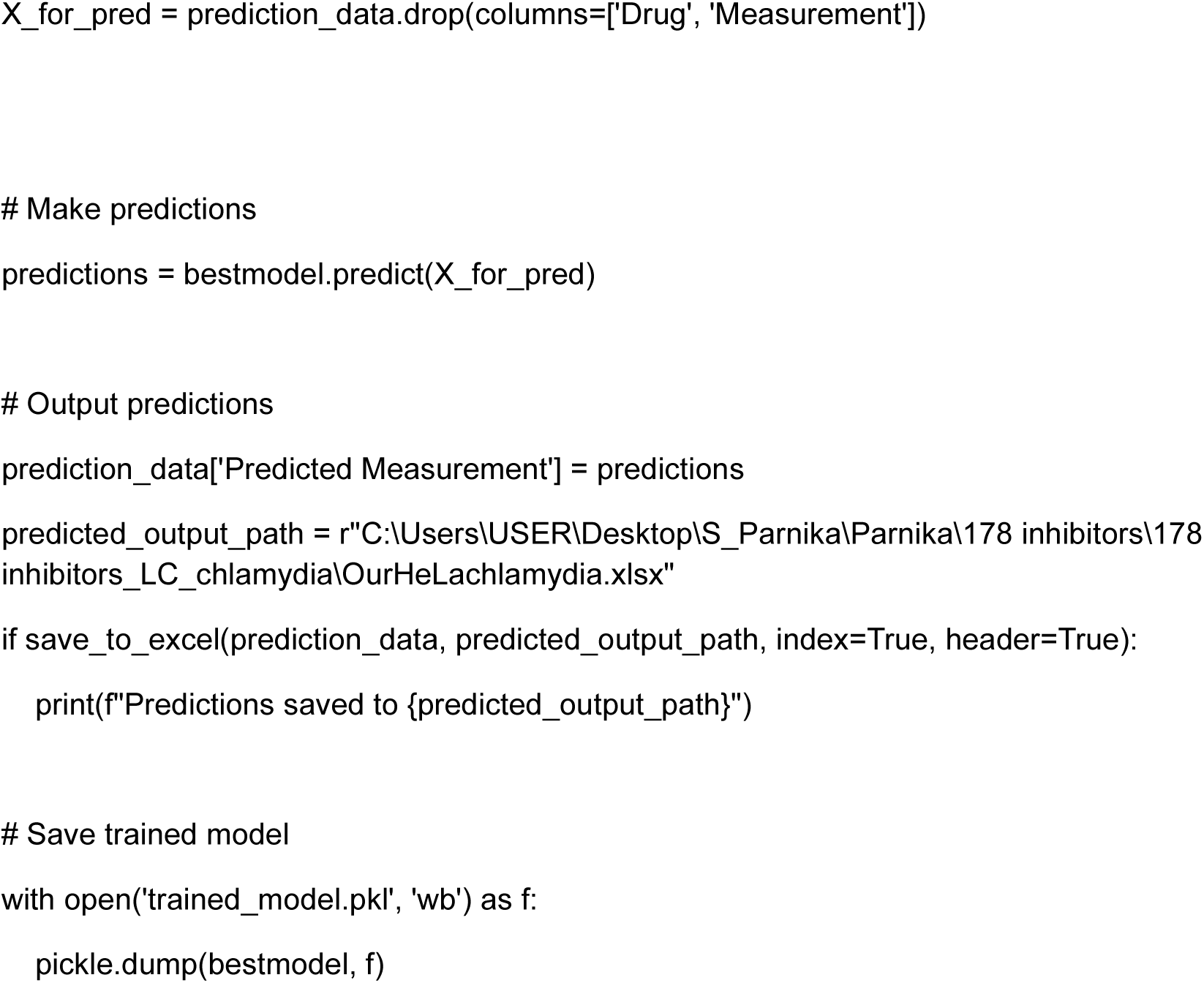

## References

1. Abu Kwaik, Y., and Bumann, D. (2015). Host delivery of favorite meals for intracellular pathogens. PLoS Pathog. 11, e1004866.

2. Eisenreich, W., Heesemann, J., Rudel, T., and Goebel, W. (2015). Metabolic Adaptations of Intracellullar Bacterial Pathogens and their Mammalian Host Cells during Infection (“Pathometabolism”). Microbiol. Spectr. 3, 27–58.

3. Ray, K., Marteyn, B., Sansonetti, P.J., and Tang, C.M. (2009). Life on the inside: the intracellular lifestyle of cytosolic bacteria. Nat. Rev. Microbiol. 7, 333–340.

4. Best, A., and Abu Kwaik, Y. (2019). Nutrition and bipartite metabolism of intracellular pathogens. Trends Microbiol. 27, 550–561.

5. Voth, D.E., and Heinzen, R.A. (2007). Lounging in a lysosome: the intracellular lifestyle of Coxiella burnetii. Cell. Microbiol. 9, 829–840.

6. Heinzen, R.A., Scidmore, M.A., Rockey, D.D., and Hackstadt, T. (1996). Differential interaction with endocytic and exocytic pathways distinguish parasitophorous vacuoles of Coxiella burnetii and Chlamydia trachomatis. Infect. Immun. 64, 796–809.

7. Elwell, C., Mirrashidi, K., and Engel, J. (2016). Chlamydia cell biology and pathogenesis. Nat. Rev. Microbiol. 14, 385–400.

8. Hackstadt, T., Scidmore, M.A., and Rockey, D.D. (1995). Lipid metabolism in Chlamydia trachomatis-infected cells: directed trafficking of Golgi-derived sphingolipids to the chlamydial inclusion. Proc. Natl. Acad. Sci. USA 92, 4877–4881.

9. Steele-Mortimer, O. (2008). The Salmonella-containing vacuole: moving with the times. Curr. Opin. Microbiol. 11, 38–45.

10. Macdonald, L.J., Graham, J.G., Kurten, R.C., and Voth, D.E. (2014). Coxiella burnetii exploits host cAMP-dependent protein kinase signalling to promote macrophage survival. Cell. Microbiol. 16, 146–159.

11. Voth, D.E., and Heinzen, R.A. (2009). Sustained activation of Akt and Erk1/2 is required for Coxiella burnetii antiapoptotic activity. Infect. Immun. 77, 205–213.

12. Hackstadt, T., Scidmore-Carlson, M.A., Shaw, E.I., and Fischer, E.R. (1999). The Chlamydia trachomatis IncA protein is required for homotypic vesicle fusion. Cell. Microbiol. 1, 119–130.

13. Sah, P., Nelson, N.H., Shaw, J.H., and Lutter, E.I. (2019). Chlamydia trachomatis recruits protein kinase C during infection. Pathog. Dis. 77, ftz061.

14. Hussain, S.K., Broederdorf, L.J., Sharma, U.M., and Voth, D.E. (2010). Host Kinase Activity is Required for Coxiella burnetii Parasitophorous Vacuole Formation. Front. Microbiol. 1, 137.

15. Sah, P., and Lutter, E.I. (2020). Hijacking and use of host kinases by chlamydiae. Pathogens 9, E1034.

16. Knight, Z.A., Lin, H., and Shokat, K.M. (2010). Targeting the cancer kinome through polypharmacology. Nat. Rev. Cancer 10, 130–137.

17. Gujral, T.S., Peshkin, L., and Kirschner, M.W. (2014). Exploiting polypharmacology for drug target deconvolution. Proc. Natl. Acad. Sci. U. S. A. 111, 5048–5053.

18. Olson, A.T., Kang, Y., Ladha, A.M., Zhu, S., Lim, C.B., Nabet, B., Lagunoff, M., Gujral, T.S., and Geballe, A.P. (2023). Polypharmacology-based kinome screen identifies new regulators of KSHV reactivation. PLoS Pathog. 19, e1011169.

19. Arang, N., Kain, H.S., Glennon, E.K., Bello, T., Dudgeon, D.R., Walter, E.N.F., Gujral, T.S., and Kaushansky, A. (2017). Identifying host regulators and inhibitors of liver stage malaria infection using kinase activity profiles. Nat. Commun. 8, 1232.

20. Dankwa, S., Dols, M.-M., Wei, L., Glennon, E.K.K., Kain, H.S., Kaushansky, A., and Smith, J.D. (2021). Exploiting polypharmacology to dissect host kinases and kinase inhibitors that modulate endothelial barrier integrity. Cell Chem. Biol. 28, 1679–1692.e4.

21. Wei, L., Barrie, U., Aloisio, G.M., Khuong, F.T.H., Arang, N., Datta, A., Kaushansky, A., and Wetzel, D.M. (2024). Using machine learning to dissect host kinases required for Leishmania internalization and development. bioRxivorg. 10.1101/2024.05.16.593986.

22. Ding, Y., Peng, Q., Song, Z., and Chen, H. (2023). Variable selection and regularization via arbitrary rectangle-range generalized elastic net. Stat. Comput. 33. 10.1007/s11222-023-10240-4.

23. McDonough, J.A., Newton, H.J., Klum, S., Swiss, R., Agaisse, H., and Roy, C.R. (2013). Host pathways important for Coxiella burnetii infection revealed by genome-wide RNA interference screening. MBio 4, e00606–e00612.

24. Jimmidi, R., Monsivais, D., Ta, H.M., Sharma, K.L., Bohren, K.M., Chamakuri, S., Liao, Z., Li, F., Hakenjos, J.M., Li, J.-Y., et al. (2024). Discovery of highly potent and ALK2/ALK1 selective kinase inhibitors using DNA-encoded chemistry technology. Proc. Natl. Acad. Sci. U. S. A. 121, e2413108121.

25. Glennon, E.K.K., Wei, L., Roobsoong, W., Primavera, V.I., van Zyl, E.M., Tongogara, T., Yee, C.B., Sattabongkot, J., and Kaushansky, A. (2026). Host kinase regulation of Plasmodium vivax dormant and replicating liver stages. PLoS Negl. Trop. Dis. 20, e0014053.

26. Ren, Q., Robertson, S.J., Howe, D., Barrows, L.F., and Heinzen, R.A. (2003). Comparative DNA microarray analysis of host cell transcriptional responses to infection by Coxiella burnetii or Chlamydia trachomatis. Ann. N. Y. Acad. Sci. 990, 701–713.

27. Ruberto, A.A., Maher, S.P., Vantaux, A., Joyner, C.J., Bourke, C., Balan, B., Jex, A., Mueller, I., Witkowski, B., and Kyle, D.E. (2022). Single-cell RNA profiling of Plasmodium vivax-infected hepatocytes reveals parasite- and host-specific transcriptomic signatures and therapeutic targets. Front. Cell. Infect. Microbiol. 12, 986314.

28. Wang, S.-S., Zhou, C.-X., Elsheikha, H.M., He, J.-J., Zou, F.-C., Zheng, W.-B., Zhu, X.-Q., and Zhao, G.-H. (2022). Temporal transcriptomic changes in long non-coding RNAs and messenger RNAs involved in the host immune and metabolic response during Toxoplasma gondii lytic cycle. Parasit. Vectors 15, 22.

29. Blanco-Melo, D., Nilsson-Payant, B.E., Liu, W.-C., Uhl, S., Hoagland, D., Møller, R., Jordan, T.X., Oishi, K., Panis, M., Sachs, D., et al. (2020). Imbalanced host response to SARS-CoV-2 drives development of COVID-19. Cell 181, 1036–1045.e9.

30. Xu, M., Wang, X., Li, Y., Geng, X., Jia, X., Zhang, L., and Yang, H. (2021). Arachidonic acid metabolism controls macrophage alternative activation through regulating oxidative phosphorylation in a PPARγ-dependent manner. Front. Immunol. 12, 618501.

31. Liang, H., Benard, O., Kumar, V., Griffen, A., Ren, Z., Sivalingam, K., Wang, J., de Simone Benito, E., Zhang, X., Zhang, J., et al. (2025). Wnt/ERK/CDK4/6 activation in the partial EMT state coordinates mammary cancer stemness with self-renewal and inhibition of differentiation. Br. J. Cancer 133, 986–1002.

32. Bone Morphogenetic Proteins. Cold Spring Harbor Perspectives in Biology 8.

33. Settembre, C., Di Malta, C., Polito, V.A., Garcia Arencibia, M., Vetrini, F., Erdin, S., Erdin, S.U., Huynh, T., Medina, D., Colella, P., et al. (2011). TFEB links autophagy to lysosomal biogenesis. Science 332, 1429–1433.

